# How the *Drosophila* Cryptochrome C-terminus mediates magnetosensitivity

**DOI:** 10.1101/2025.09.15.676315

**Authors:** Adam A Bradlaugh, Anna L Munro, Daniel Kattnig, Moona Kurttila, Noboru Ikeya, Alex Hoose, Sanjai Patel, Susanna Campesan, Charalambos P. Kyriacou, Ezio Rosato, Jonathan R. Woodward, Alex R. Jones, Richard A. Baines

**Affiliations:** Division of Neuroscience, School of Biological Sciences, Faculty of Biology, Medicine and Health, University of Manchester, Manchester Academic Health Science Centre, Manchester, M13 9PL, UK; Living Systems Institute, University of Exeter, Exeter, Devon, EX4 4QD, UK; Department of Physics, University of Exeter, Stocker Rd, Exeter EX4 4QL; Biometrology, Chemical and Biological Sciences Department, National Physical Laboratory, Teddington, Middlesex, TW11 0LW, UK; Graduate School of Arts and Sciences, The University of Tokyo, 3-8-1 Komaba, Meguro-ku, Tokyo 153-8902, Japan; Manchester Fly Facility, Faculty of Biology, Medicine and Health, University of Manchester, M13 9PT, UK; Division of Genetics and Genome Biology, School of Biological and Biomedical Sciences, College of life Sciences, University of Leicester, LE1 7RH

## Abstract

The Earth’s magnetic field plays an important role in the seasonal migrations of many species of animals. A Cryptochrome (CRY)-based radical pair mechanism (RPM) has been suggested to underlie the mechanistic basis of animal magnetosensitivity and navigation. The quantum spin state of a radical pair involving flavin adenine dinucleotide (FAD) bound to CRY in the canonical pocket is sensitive to external magnetic fields that can alter the signalling concentration of activated CRY^1–5^. However, several experimental observations challenge this model including the finding that the C-terminal fragment of *Drosophila* CRY (*Dm*CRY), which lacks any canonical FAD binding pocket, and human CRY2, which lacks affinity for FAD, are sufficient to support magnetosensitivity^6–9^. Here, we use all-atom molecular dynamic (MD) simulations, alongside *in vitro* and *in vivo* analyses to reveal that the C-terminus of *Drosophila* CRY (*Dm*CRY-CT) binds FAD. FAD binding is required for transduction of a magnetic signal within cells, and, *in vitro*, initiates formation of high molecular weight *Dm*CRY-CT oligomers, including large insoluble aggregates reminiscent of CRY photobodies observed in plants^10–14^. These results provide a plausible mechanistic basis for several experimental observations that have reported non-canonical magnetosensitivity in animals.

## Introduction

Many animals sense the Earth’s geomagnetic field^15^. Whilst well-established, the identity of the magnetoreceptor and the molecular basis of magnetic signal transduction remains controversial. The Cryptochrome (CRY)-based radical pair mechanism (RPM) has gained traction in recent years. In brief, absorption of blue light (BL) by flavin adenine dinucleotide (FAD), bound to CRY, initiates a series of cascaded electron transfers along a chain of tryptophan (Trp) residues towards FAD to form a spin-correlated, FAD^•−^ / Trp^•+^ radical pair (RP)^1,3–5,16,17^. Formation of the FAD radical correlates with the active state of CRY; for example, in *Drosophila melanogaster* the radical triggers movement of the *Dm*CRY C-terminal tail (CTT), which allows CRY to interact with downstream effectors^18–25^. The current CRY-dependent model of RP magnetoreception proposes that the relative populations of spin states of the RP are altered by a magnetic field (MF). The differential reactivity of the singlet and triplet spin states, in which the former can recombine back to the inactive state, while the latter cannot, means that the MF modulates the population of the RP and hence the concentration of active CRY^1,3–5,17^. This canonical RP model mediated by FAD^•−^/ Trp^•+^ has been questioned^6,8,26–29^.

A striking observation that challenges the canonical FAD-Trp RP model is that the expression of just the C-terminal 52 amino acids of *Drosophila* CRY (*Dm*CRY-CT) is sufficient to mediate magnetosensitivity in both a *Drosophila* adult circadian and a larval neuronal assay^6–8,30^. The *Dm*CRY-CT fragment lacks the N-terminal FAD-binding pocket and the tetrad of Trp residues critical for the electron transfer chain. Additionally, increasing intracellular FAD concentration to non-physiological levels in the fly neuronal assay is sufficient to support non-CRY-dependent magnetosensitivity^8^. The C-terminus of *Dm*CRY contains putatively characterised PDZ-interaction motifs and a calmodulin-binding domain^24,25,31^ which may facilitate the targeting of a FAD bound *Dm*CRY-CT complex to neuronal scaffolds such as the protein ‘inactivation no afterpotential D’ (INAD), as reported in the *Drosophila* visual system^25^, or with plasma membrane-bound ion channels such as the Shaker K^+^ channel^32,33^. These downstream scaffolds could transfer the RP quantum response into behavioural output. While free FAD is able to support magnetic field effects (MFEs) via an intramolecular RP (Extended Data Fig. 1A), sizeable effects require the adoption of an “open configuration” which is not favoured at neutral pH^34,35^.

In this study, we report that FAD and the *Dm*CRY-CT bind. Molecular dynamic (MD) simulations identify electrostatically bound configurations with highly favourable interaction energies. We further show binding between *Dm*CRY-CT and FAD *in vitro*, in a process that drives the formation of peptide oligomers in a light-sensitive manner. Point mutations in *Dm*CRY-CT, predicted as essential for CT-FAD complex formation, abolish the capability of *Dm*CRY-CT to support magnetosensitivity. Additionally, we show that the PDZ-interaction motifs in *Dm*CRY-CT are required to mediate magnetosensitivity within a neuron.

## Results and Discussion

### Molecular Dynamic Simulations show FAD binds *Dm*CRY-CT

In previous work^6,8^, we reported that *Dm*CRY-CT is sufficient to support magnetosensitivity in neurons. To do so, we considered that *Dm*CRY-CT might bind FAD and that binding configurations exist that stabilize FAD’s open-configuration, which shows larger MFEs because of longer RP lifetimes and more favourable relative energies between the different spin-states^34^. We employed MD simulations to test these hypotheses.

We excised the *Dm*CRY C-terminal tail (CT) from the published crystal structure (PDB ID: 4GU5; net charge of CT: +1 e) and extended it by the missing three valine residues (540– 542, Extended Data Fig. 1B). Extensive MD simulations (NpT ensemble; 298 K, 1 atm) yielded an equilibrated solution structure of the free *Dm*CRY-CT peptide. In this structure, the two α-helices (residues 498-516 and 528-535) and the connecting loop form a U-shaped structure. To identify binding configurations, we performed short-time (≤ 30 ns) MD simulations after introducing a FAD molecule (in its closed configuration) into the solution, positioned approximately 1 nm from the peptide in various relative orientations. The FAD was consistently captured by the peptide, typically during the equilibration phase. Several binding configurations are possible, but two representative configurations were identified from these simulations where FAD binds in an open conformation as hypothesised (Fig. 1A). These were selected for more detailed analysis using extended MD simulations.

**Figure 1.**
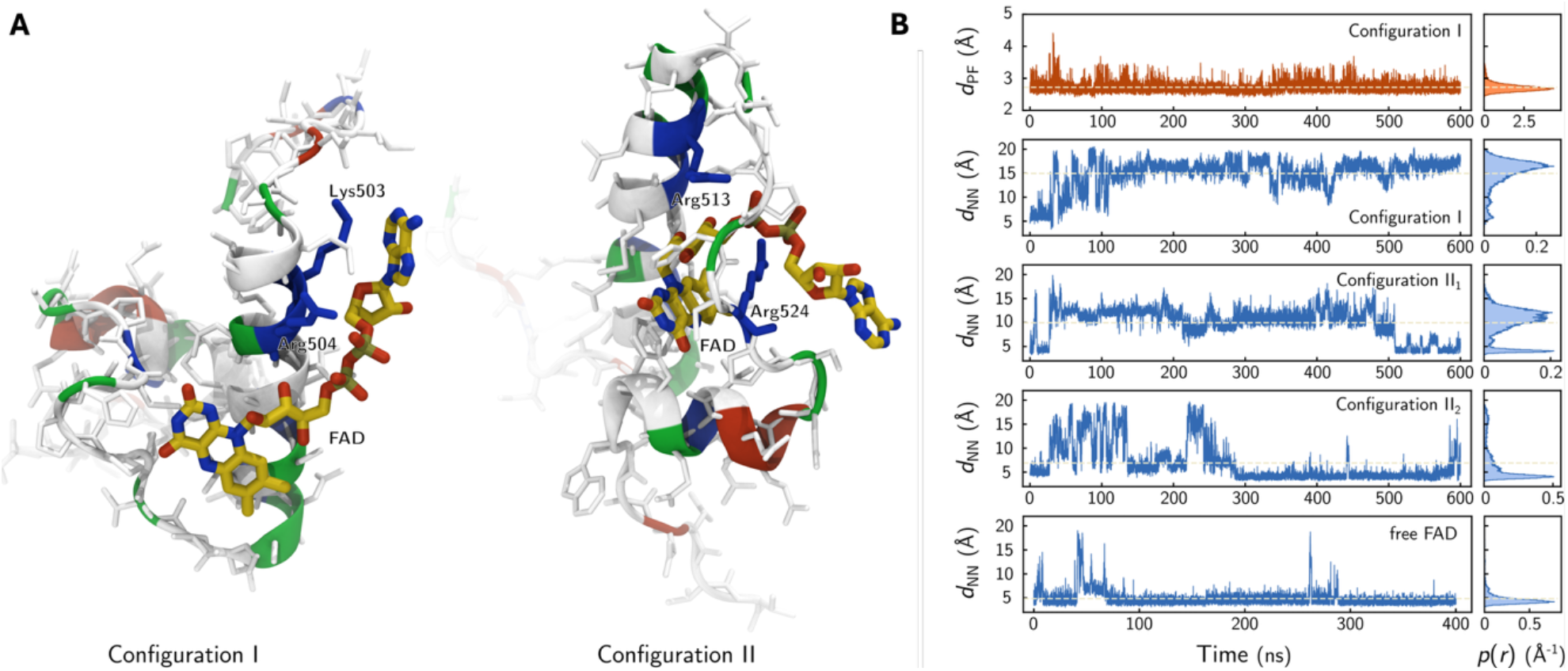
Molecular Dynamic Simulations suggest electrostatic binding of FAD directly to *Dm*CRY-CT, with two example configurations promoting the adoption of an ‘open’ conformation in FAD. A) Snapshots of FAD bound to the *Dm*CRY-CT in *configuration I* (left; binding Arg504 and Lys503) and *configuration II* (right; binding Arg524 and Arg513). B) Time evolution of the peptide-FAD distance _*d*PF_ (red) and the inter-moity distance _*d*NN_ in FAD (blue) over selected MD trajectories. From top to bottom, the subplots represent _*d*PF_ and _*d*NN_ for FAD bound in *configuration I*; two replicas of _*d*NN_ for FAD bound in *configuration II* (*II*1 and *II*2, respectively); and _*d*NN_ for free FAD. The left panels give the distance as a function of the simulation time after equilibration, and the right panels provide the associated probability densities. The dashed grey lines represent the mean distances. _*d*PF_ is the shorted distance between heavy atoms; _*d*NN_ corresponds to the distance of N10 in the isoalloxazine ring and N1Z in the adenine ring. _*d*PF_ shows comparable behaviour in all bound configurations considered here.

The two binding configurations between *Dm*CRY-CT and open forms of FAD exhibit FAD-peptide distances around 2.6 Å, with strong electrostatic interactions between the negatively charged diphosphate group in FAD and positively charged amino acids. These configurations showed no tendency for dissociation during extended MD simulations. In *configuration I*, FAD binds at the proximal end of the peptide, primarily through electrostatic interactions mediated by Lys503 and Arg504. In *configuration II*, FAD is partially intercalated into a groove formed by the α-helices and their connecting loop, with binding facilitated by Arg524 and Arg513. Binding energies ranged from −170 kcal/mol for *configuration I* to −280 kcal/mol for *configuration II* (with partial intercalation); protein-FAD distances reflect van der Waals contact (*d*_*PF*_ in Table 1). The interaction was dominated by electrostatic forces, with van der Waals contributions accounting for a smaller fraction (−30 to −50 kcal/mol, Table 1). The ‘open’ state of FAD in each configuration is reflected in the increased distance between the N10 atom in the isoalloxazine ring and the N1Z atom in the adenine ring (Fig. 1B). In *configuration I*, this distance expanded to 15.0 ± 3.1 Å from 4.8 ± 1.7 Å in free FAD. In *configuration II*, it increased to 9.9 ± 3.1 Å and 6.9 ± 4.0 Å in two replicate runs, suggesting it might be less well-suited to magnetic sensing. The FAD-radius of gyration shows the same trends. Key parameters for the *Dm*CRY-CT/FAD complexes, including peptide-FAD distances, binding energies, interaction sites, and inter-moiety distances within FAD are reported in Table 1.

**Table 1.** FAD binding behaviour as observed during production MD simulation runs for the wildtype peptide with FAD initially bound in *configurations I, II*_***1***_, **and *II***_***2***_.

| **Configuration** | **$t_{\text{sim}}$**<br>**(ns)** | **$d_{PF}$**<br>**(Å)** | **$d_{NN}$**<br>**(Å)** | **Main**<br>**Interactors** | **$E_b$**<br>**(kcal/mol)** | **$E_{es}$**<br>**(kcal/mol)** | **$E_{vdV}$**<br>**(kcal/mol)** |
| --- | --- | --- | --- | --- | --- | --- | --- |
| **I** | 600 | 2.72±0.15 | 15.0±3.1 | 503, 504, 510 | -172±80 | -138±79 | -34±8 |
| **II<sub>1</sub>** | 600 | 2.63±0.07 | 6.9±4.0 | 524, 513, 510 | -277±55 | -249±55 | -28±7 |
| **II<sub>2</sub>** | 500 | 2.67±0.12 | 9.9±3.1 | 524, 513, 510 | -252±74 | -206±73 | -46±9 |

$t_{\text{sim}}$ : simulation time; $d_{PF}$ : smallest distance between the peptide and FAD, measured for non-hydrogen atoms; $d_{NN}$ : distance of N10 in the isoalloxazine moiety and N1Z in the adenine moiety of FAD; main interactors: the three residues of most negative binding energy; $E_b$ , $E_{es}$ , and $E_{vdV}$ : the total, electrostatic, and van der Waals binding energy averaged over the MD run.

### *In vitro* analysis confirms *Dm*CRY-CT/FAD binding and reveals photobody formation

Supporting MD simulations, we found strong evidence for the formation of *Dm*CRY-CT/FAD complexes *in vitro*. To aid with solubility, we used a synthesised variant of the *Dm*CRY-CT peptide, where the C-terminal DVVV is removed (similar to the truncation of full-length *Dm*CRY to achieve its crystal structure^22,23^) (Extended Data Fig. 1B). Although this region represents a putative PDZ binding motif, whose presence is required in the single neuron assay (see below), there is no evidence from MD simulations that these residues are involved in interactions that stabilise FAD binding. Binding of FAD to *Dm*CRY-CT was first investigated using size exclusion chromatography (SEC), measuring elution volume of eluting species by absorption at 280 nm. In general, larger species elute from a SEC column at a smaller elution volume. Solutions of the isolated *Dm*CRY-CT peptide (which is larger) and FAD eluted with overlapping peaks at 17.3 mL and 18.0 mL, respectively (Extended Data Fig. 3A). Solutions of the peptide with increasing ratios of FAD elute with a shoulder at 17.0 mL to the peptide elution peak, which increases in magnitude with increasing FAD concentration (Fig. 2A and Extended Data Fig. 3B). This is consistent with the formation of soluble complexes driven by the addition of FAD and are indicative of binding between *Dm*CRY-CT and FAD. There is also evidence for the formation of much larger complexes during the titration. First, a SEC peak elutes at 8.2 mL (Fig. 2A), consistent with soluble oligomers that are reminiscent of CRY photobodies, which appear to have a signalling role in plants^11–14^. Second, UV-visible spectra of different *Dm*CRY-CT/FAD ratio mixtures, normalised to the FAD concentration, show a baseline sloping upwards from longer to shorter wavelengths (Fig. 2B). This baseline, whose magnitude is enhanced with increasing proportion of FAD, is consistent with higher molecular weight complexes, in the form of insoluble peptide aggregates, that scatter incident light. Centrifugation to remove the precipitate also removes the sloping baseline (*i.e*., removes scattering) but leaves the concentration of FAD virtually unchanged (Extended Data Fig. 3C). This suggests that if higher order complex formation is driven by FAD binding, only a very small number of FAD molecules are required to ‘seed’ aggregation of the *Dm*CRY-CT peptide. These large, insoluble complexes are possibly not present in the SEC samples due to sample filtering prior to loading. The formation of the soluble complexes is further enhanced following illumination with blue light (450 nm, Fig. 2C), where the shoulder to the peptide peak now forms a discrete peak, or when the solutions were prepared and SEC run under ambient light (Extended Data Fig. 3D).

**Figure 2.**
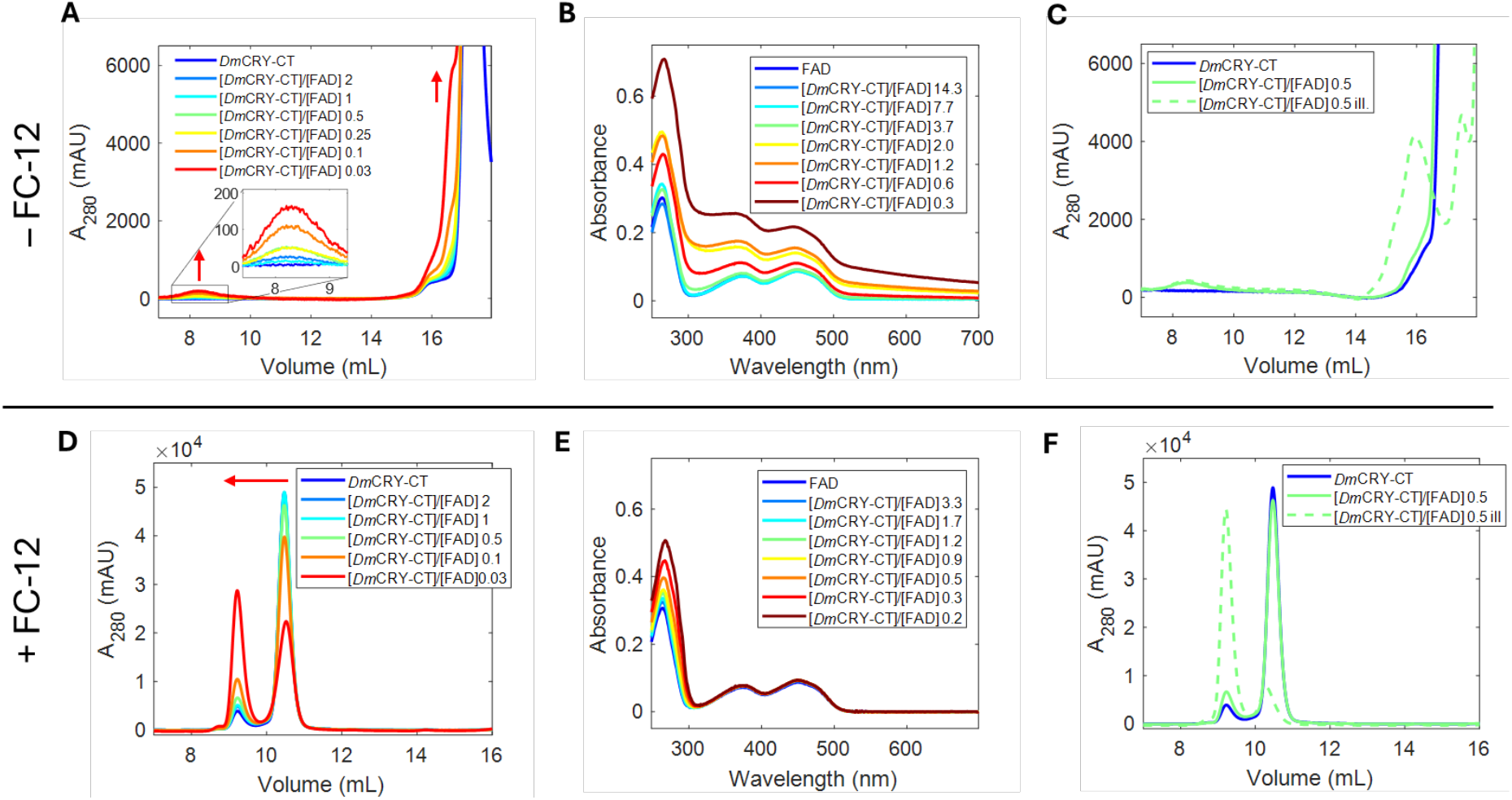
SEC elution data and UV-visible absorption spectra from *Dm*CRY-CT/FAD titrations in the absence of the detergent FC-12. A), a 30 μM solution of the *Dm*CRY-CT peptide is added to the column with increasing proportions of FAD. The figure legends display the molar ratio between *Dm*CRY-CT and FAD and the peak at elution volume of ∼ 8.2 mL is expanded within the inset for clarity. B) spectra acquired from a 7.3 μM solution of FAD with increasing proportions of the *Dm*CRY-CT peptide. C) SEC elution of the *Dm*CRY-CT peptide (blue) is compared to elutions for a fixed *Dm*CRY-CT / ratio either prepared and run in darkness (green) or following illumination with blue light (green dashed). D-F) Equivalent data to (A-C) but for solutions prepared in the presence of FC-12. FAD elutes at 18.0 mL. All SEC elution volumes are monitored using absorbance at 280 nm and solutions are prepared and run in darkness unless stated otherwise. See panel legends for *Dm*CRY-CT/FAD ratios.

Adding the detergent Fos-choline 12 (FC-12) was sufficient to prevent the formation of *Dm*CRY-CT/FAD oligomers and higher-order aggregates. The *Dm*CRY-CT elution volume changes significantly in the presence of FC-12 (10.5 mL, Fig. 2D and Extended Data Fig. 3E), because association of FC-12 to the peptide forms larger species. The addition of FAD still leads to the formation of small, soluble complexes in the presence of FC-12. Because the peptide peak no longer overlaps with that of free FAD, complex formation is clearly observed: the elution peak from peptide alone (10.5 mL) diminishes as the peak from the complex that replaces it (9.2 mL) grows in. There is no evidence of a separate elution peak owing to soluble oligomers in the presence of FC-12, although there’s a chance the peptide / FC-12 elution volume might obscure this. That said, the UV-visible spectra of the different *Dm*CRY-CT/FAD mixtures show no evidence of a sloping baseline in the presence of FC-12 (Fig. 2E), so we tentatively conclude that higher order structures (both soluble and insoluble) are not formed. Again, the yield of small, soluble, complexes increase significantly following illumination (Fig. 2F and Extended Data Fig. 3F).

To investigate further the formation of higher order complexes, we conducted light microscopic measurements on *in vitro* FAD / *Dm*CRY-CT mixtures in the presence and absence of FC-12. Sample mixtures of 100 μM *Dm*CRY-CT with varying FAD concentrations were prepared and imaged under irradiation with 450 nm laser light (Fig. 3A). Samples containing FAD of concentrations at, or greater than, 1 μM formed fluorescent particles. Aggregate formation was completely supressed in the presence of FC-12 (Fig. 3B). Moreover, adding FC-12 after particle formation produced a sample where the particles were no longer observed; presumably having redissolved in the detergent solution (Extended Data Fig. 4A-C). The emissive particle formation was observed immediately upon imaging regardless of whether the sample was prepared in dark or under room light. Irradiation (450 nm), however, leads to the accumulation of increasing numbers and size of particles in and around the irradiation spot (Fig. 3C, Extended Data Fig. 5A, video available at Harvard Dataverse: https://doi.org/10.7910/DVN/GLW1UV). A non-irradiated sample left for the same period (3 hours) does not show this increase (Extended Data Fig. 5B).

**Figure 3.**
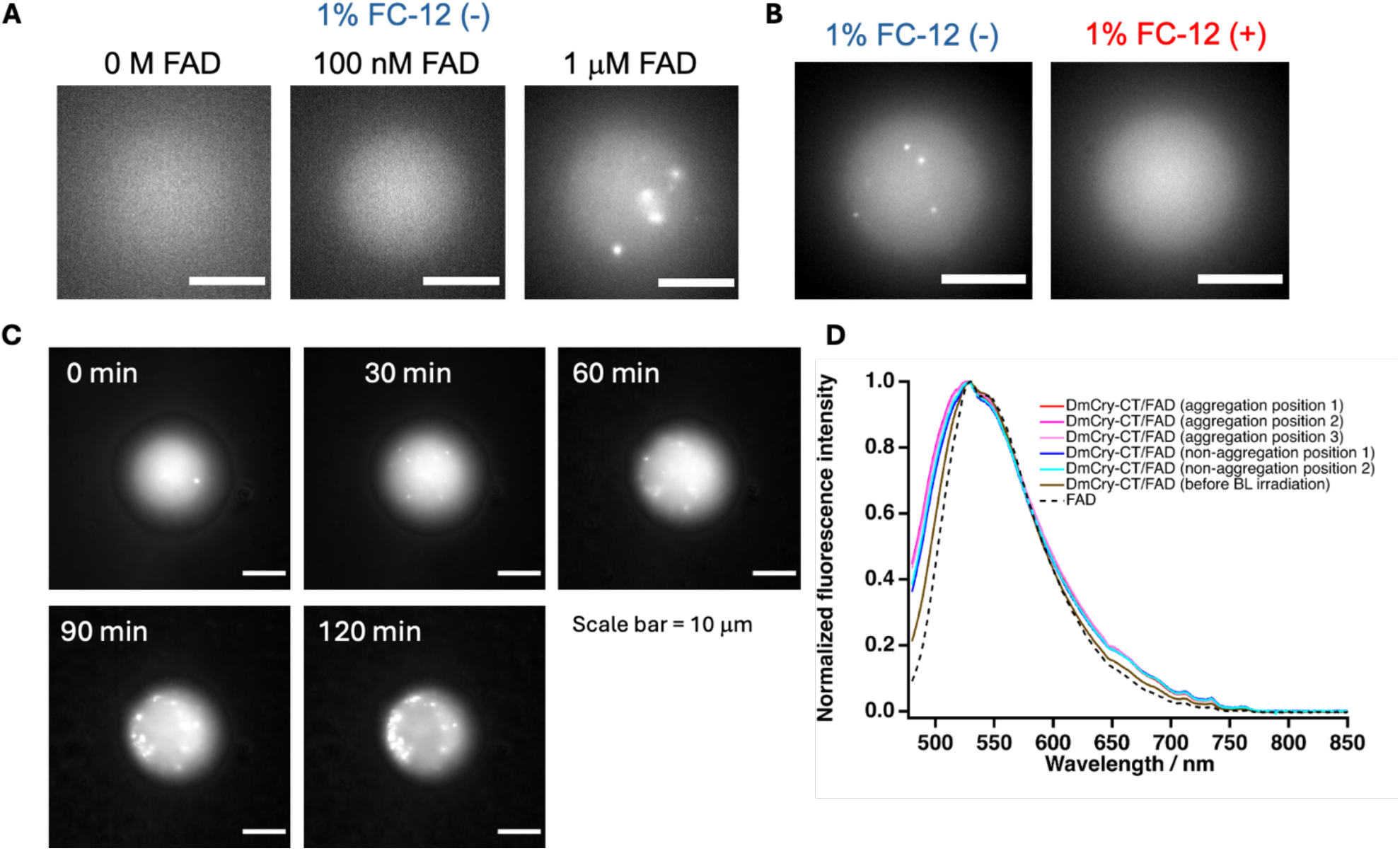
Fluorescence microspectroscopy-based *in vitro* characterisation of *Dm*CRY-CT/FAD binding and aggregation. A) Representative fluorescence images of 100 μM *Dm*CRY-CT solution (pH 7.4) at varying FAD concentrations in the absence of detergent. B) Representative fluorescence images of 100 μM *Dm*CRY-CT solution (pH 7.4) with 10 μM FAD in the presence and absence of detergent (1 % FC-12). C) Fluorescence images showing aggregate formation in the *Dm*CRY-CT (100 μM) and FAD (10 μM) mixture (pH 7.4) under continuous irradiation at 450 nm on a long timescale (Excitation intensity = 2.1 kW/cm^2^). D) Fluorescence spectra of 100 μM *Dm*CRY-CT and 10 μM FAD at aggregation and non-aggregation positions after long time 450 nm irradiation, and at the non-aggregation position before irradiation. Each spectrum is normalized to its maximum intensity.

To identify the biomolecular origin of the particle fluorescence, we recorded fluorescence spectra from both aggregation and non-aggregation positions (Fig. 3D). The spectra from both were similar and reflect a characteristic FAD spectrum. Furthermore, in the irradiated *Dm*CRY-CT/FAD sample, the spectra change shape compared to free FAD and non-irradiated *Dm*CRY-CT/FAD: there is a slightly higher intensity between 480 nm and 530 nm relative to longer wavelengths, which is consistent with FAD binding and the light-induced enhancement of this binding^36,37^. We therefore conclude that FAD is present within the particles formed and that there are no other significant sources of fluorescence from the particles, which is consistent with FAD-binding to *Dm*CRY-CT and that this binding drives peptide oligomerisation / aggregation.

Our *in vitro* analysis provides evidence for the formation of at least three populations of *Dm*CRY-CT/FAD complex: i) small, soluble complexes; ii) larger, soluble oligomers; and iii) larger, insoluble aggregates. Although the formation of these complexes doesn’t appear to require light, in each case the yield is enhanced by illumination with BL. *In vitro* fluorescence quenching experiments (Extended Data Fig. 6-8) suggest that FAD binds to the *Dm*CRY peptide in different ways. First, in a non-emissive confirmation that forms the basis of complexes that remain small and soluble; and second, in a confirmation that drives the formation of higher order structures (soluble oligomers and insoluble aggregates) where only a small proportion of FAD is required and the photophysical properties of FAD remain largely unchanged.

### *In vivo* electrophysiology of *Dm*CRY-CT/FAD binding mutants validates their necessity for magnetosensitivity

MD simulations were used to identify key residues required for favourable FAD binding. As described in the Extended Data, selected mutants, summarized in Table 2, showed strongly reduced binding to the protein, leading to FAD-detachment, and/or failed to support an open FAD configuration. Based on these *in silico* explorations (Fig. 1 and Extended Data Fig. 2), we made transgenic constructs for expression in our single neuron assay (Table 2).

**Table 2.** FAD binding as observed during production MDS runs for various mutants with FAD initially bound in *configurations I, II*_***1***_, **and *II***_***2***_.

| **Configuration** | **$t_{sim}$**<br>**(ns)** | **$d_{PF}$**<br>**(Å)** | **$d_{NN}$**<br>**(Å)** | **Main**<br>**Interactors** | **$E_b$**<br>**(kcal/mol)** | **$E_{es}$**<br>**(kcal/mol)** | **$E_{vdW}$**<br>**(kcal/mol)** |
| --- | --- | --- | --- | --- | --- | --- | --- |
| **I-R504E** | 400 | 5.23±6.85 | 11.2±5.8 | 503, 510, 526 | -119±77 | -94±76 | -26±8 |
| **II<sub>1</sub>-R524K** | 200 | 2.67±0.08 | 4.4±0.6 | 513, 510, 524 | -231±62 | -202±62 | -28±4 |
| **II<sub>1</sub>-R524A** | 200 | 2.69±0.11 | 4.3±0.9 | 513, 510, 506 | -155±32 | -126±32 | -29±4 |
| **II<sub>1</sub>-R524E** | 400 | 2.69±0.10 | 17.7±3.4 | 513, 510, 527 | -121±26 | -87±25 | -35±4 |
| **II<sub>2</sub>-R524E** | 400 | 2.73±0.27 | 7.4±2.7 | 504, 505, 510 | -126±64 | -96±63 | -30±7 |

$t_{sim}$ : simulation time; $d_{PF}$ : smallest distance between the peptide and FAD, measured for non-hydrogen atoms; $d_{NN}$ : distance of N10 in the isoalloxazine moiety and N1Z in the adenine moiety of FAD; main interactors: the three residues of most negative binding energy; $E_b$ , $E_{es}$ , and $E_{vdW}$ : the total, electrostatic, and van der Waals binding energy averaged over the MD run.

We have previously reported that expression of *Dm*CRY-CT in a larval neuron (the aCC motoneuron) is sufficient to support magnetosensitivity^8^. By contrast, expression of *Dm*CRY-CT^R504E^ (a positive for negative residue substitution), affecting binding *via configuration I*, did not support magnetosensitivity (Fig. 4A, *P* = 0.9975). Equally, expression of *Dm*CRY-CT^R524E^ (with a similar positive for negative residue substitution), required for FAD binding in *configuration II*, did not support magnetosensitivity (Fig. 4A, *P* = 0.8603). The lack of redundancy between the two configurations suggests that both are necessary for magnetosensitivity. The conservative mutation R524A (positive to neutral, *configuration I*) produced a slight but non-significant trend towards magnetosensitivity (Fig. 4A, *P* = 0.2247), underscoring the importance of charged residues to support electrostatic attraction.

**Figure 4.**
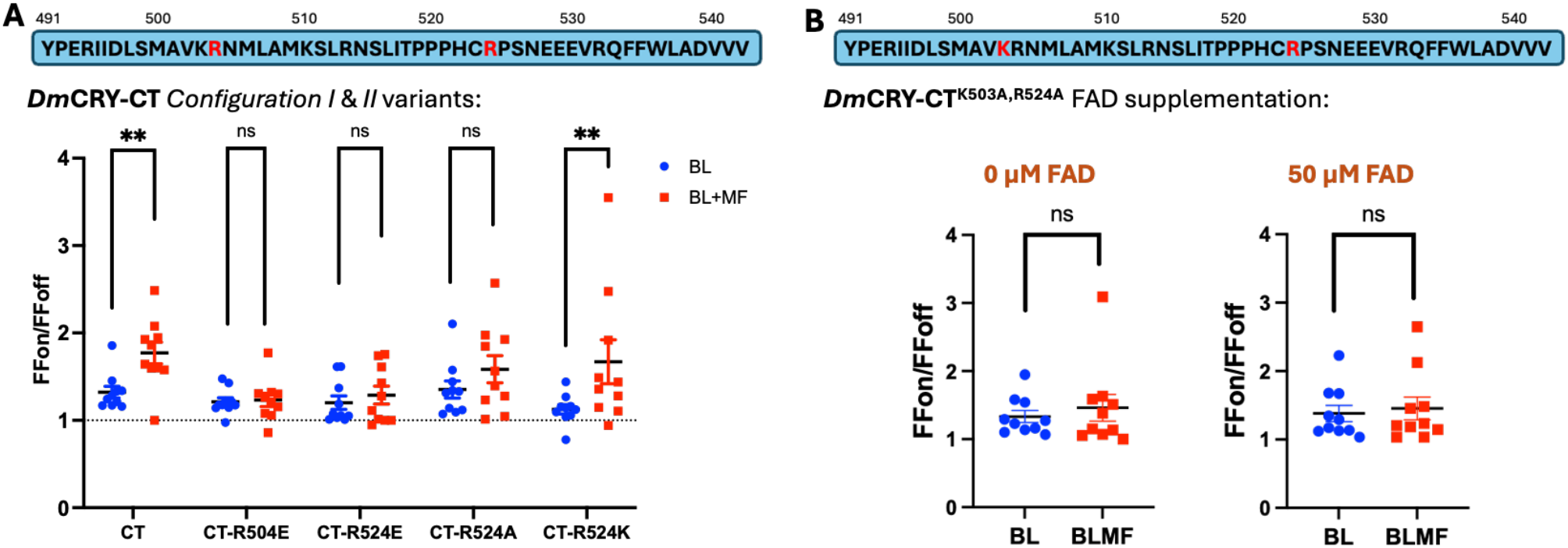
Disruption to FAD binding nullifies magnetosensitivity *in vivo*. A) The firing fold change of aCC motoneurons expressing *Dm*CRY-CT/FAD binding variants when exposed to BL with, or without, a MF (100 mT). Expression of *Dm*CRY-CT results in an increase in neuronal firing 1.32-fold (*P* = 0.0029), which is potentiated by the presence of a MF to 1.77-fold change (BL *vs* BL+MF, *P* = 0.0050). A substitution for a negative residue in *Dm*CRY-CT^R504E^ (*configuration I*) gives a modest BL effect of 1.21-fold change (*P* = 0.0041), which is not potentiated by MF exposure (1.23-fold change, BL *vs* BL+MF, *P* = 0.9975). The substitution for a negative sidechain at a residue involved in FAD binding at *configuration II, Dm*CRY-CT^R524E^, gives a modest BL effect of 1.2-fold change (*P* = 0.0280), which is also not affected by MF exposure (1.29-fold change, BL *vs* BL+MF, *P* = 0.8603). The replacement of the same residue for a neutral alanine in *Dm*CRY-CT^R524A^ also resulted in a slight BL response of 1.35-fold change (*P* = 0.0076) but no significant potentiation in a MF (1.59-fold change, BL *vs* BL+MF, *P* = 0.2247), whereas the conservation of a positive residue in *Dm*CRY-CT^R524K^, which again exhibits BL sensitivity (1.13-fold change, *P* = 0.0409), and also retains magnetosensitivity (1.67-fold change, BL *vs* BL+MF, *P* = 0.048). B) Double mutant of both configurations in which the key positive residue for each configuration is substituted for a neutral alanine. In the presence and absence of supplemented FAD (50 µM) a small but significant BL response is seen (0 µM FAD: *P* = 0.0034, 1.33-fold change, 50 µM FAD: *P* = 0.0062 1.38-fold change). No MF potentiation of this effect is seen in the absence of additional FAD (0 µM FAD, *P* = 0.5576, 1.33-*vs* 1.46-fold change), or when 50 µM FAD is supplemented to the cell (50 µM FAD, *P* = 0.7181, 1.38-*vs* 1.46-fold change). Raw action potential counts for these recordings and their controls are found in Extended Data Fig. 9 and 11. Unpaired t-tests (two-tailed) were used for BL *vs* BL+MF comparisons, and paired t-tests (two-tailed) were used to determine BL responses. n = 10 per genotype, per condition.

Expression of *Dm*CRY-CT^R524K^ (a positive-to-positive substitution, *configuration II*) was, as predicted, able to support magnetosensitivity (Fig. 4A, *P* = 0.0486).

We previously demonstrated that supplementing the internal recording saline in the patch electrode with 200 µM FAD was sufficient to confer magnetosensitivity in the absence of the *Dm*CRY-CT^8^. Supplementing with 50 µM FAD, while not sufficient to mediate MF effects alone, boosted MF effects in motoneurons expressing *Dm*CRY-CT^8^. The double mutant *Dm*CRY-CT^K503A, R524A^, consisting of conservative positive-to-neutral substitutions at positions 503 and 524, did not support magnetosensitivity (Fig. 4B, 0 µM FAD, *P* = 0.5576) even with additional 50 µM FAD (Fig. 4B, 50 µM FAD, *P* = 0.7181). Raw action potential counts for expressed *Dm*CRY-CT transgenics and their respective non-expressed controls are presented in Extended Data Fig. 9 and 11.

### PDZ motifs in *Dm*CRY-CT are required for magnetosensitivity

We have reported that mutation of one of two putative PDZ-binding domains in full-length *Dm*CRY (V531K) is sufficient to prevent transduction of a magnetic signal in aCC motoneurons^8^. This is consistent with one/both PDZ-interaction motifs (Fig. 5A) within *Dm*CRY-CT being required for targeting the *Dm*CRY-CT/FAD complexes to downstream effectors. To validate this observation, we mutated both putative PDZ domains in *Dm*CRY-CT. We confirmed that expression of *Dm*CRY-CT^V531K^ in aCC does not support magnetosensitivity (Fig. 5B, *P* = 0.8704). Similarly, mutation of the second putative PDZ domain (V541K) produced the same negative outcome (Fig. 5C, *P* = 0.7852), despite both variants remaining sensitive to light (Extended Data Fig.10-11). These results support a scenario where bound FAD is required near to effectors required for magnetosensitivity in *Drosophila* neurons. Increased levels of FAD can support magnetosensitivity in the presence of *Dm*CRY-CT but without this peptide, FAD must be increased to a substantial non-physiological degree to observe MFEs, as previously reported^8^.

**Figure 5.**
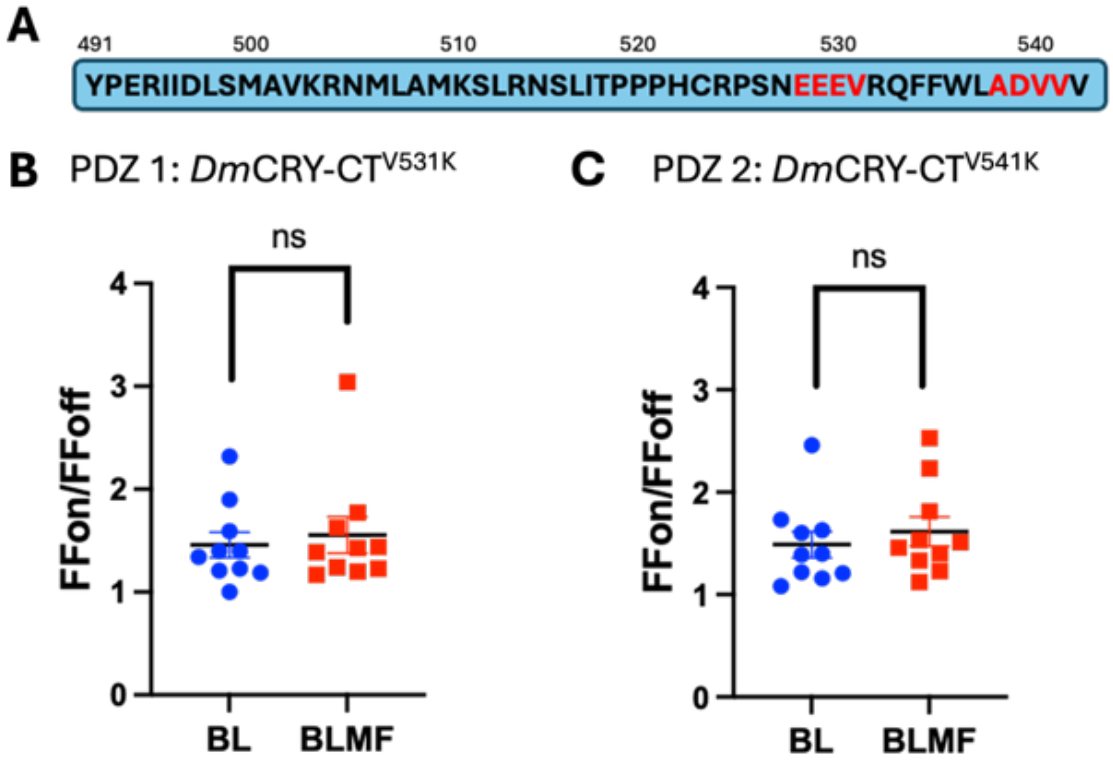
PDZ interaction motifs in the DmCRY-CT are necessary for localising the DmCRY-CT/FAD complex to downstream effectors. A) Diagram of the *Dm*CRY-CT amino acid sequence with PDZ interaction motifs highlighted in red. B) Expression of *Dm*CRY-CT^V531K^, which disrupts the PDZ motif EEEV, gave a BL response of 1.46-fold change (*P* = 0.0031) that was not modulated by MF exposure (1.55-fold change, BL *vs* BL+MF, *P* = 0.8704). C) Expression of *Dm*CRY-CT^V541K^, which disrupts the PDZ motif ADVV, gave a BL response of 1.49-fold change (*P* = 0.0031) that was not modulated by MF exposure (1.62-fold change, BL *vs* BL+MF, *P* = 0.7852). Unpaired t-tests (two-tailed) were used for BL *vs* BL+MF comparisons, and paired t-tests (two-tailed) were used to determine BL responses. n = 10 per genotype, per condition.

## Summary and Conclusions

We present *in silico* MD simulations, together with *in vitro* and *in vivo* data to show that FAD binds *Dm*CRY-CT to support MF sensing (*via* FAD) and transduction of a magnetic signal in a *Drosophila* neuron. Our simulations and mutational studies indicate the possible importance of binding configurations that can hold FAD in its open configuration, which is known to facilitate enhanced MFEs for intramolecular RPs in FAD^34,38^. Our *in vitro* analysis reveals that FAD binding to *Dm*CRY-CT results in the formation of oligomeric states, from small-soluble complexes to larger insoluble aggregates. Higher order structures have been observed previously in response to illumination for full-length CRY in plants^10–14^, with varied roles for such photobodies including degradation, signalling and transcriptional regulation. These effects are now being exploited for optogenetic control of clustering and phase transitions within cells^39^, and our data reveal the potential for magnetic modulation of such tools. Our data also suggest that light-induced oligomerisation of CRY is mediated by its C-terminus, which would be consistent with the fact that structural changes that occur upon photoactivation make available the C-terminus of CRY and the CRY-CT for interaction with partners^18–25^. However, the individual contribution of these respective higher-order structures to the observed magnetosensitivity remains to be determined. Equally important will be to determine whether FAD can bind the HsCRY2-CT, or some alternative site, other than the canonical binding pocket. HsCRY2 is sufficient to support magnetosensitivity when expressed in *Drosophila*^6,40^, even though it does not show strong FAD binding in its canonical binding pocket^9^. Finally, we confirm the vital role that two linear PDZ binding motifs play in transduction of the magnetic signal, presumably through localisation of the *Dm*CRY-CT and bound FAD to effector proteins in the cell.

## Acknowledgements

This work was supported by funding from BBSRC to RAB (BB/V005987/1) and to CPK/ER (BB/V006304/1) and a Wellcome Discovery award (311280/Z/24/Z) to RAB, CPK, ER and ARJ. We thank Wei-Hsiang Lin of the Manchester Genomics Technologies facility for generating constructs for transgenics. Work on this project benefited from the Manchester Fly Facility, established through funds from the University and the Wellcome Trust (087742/Z/08/Z). ARJ thanks the National Measurement System of the Department for Business, Energy and Industrial Strategy for funding. ER and CPK acknowledge funding from the Electromagnetic Field Biological Research Trust (BRT 15/51).

## Author Contributions

R.A.B., A.A.B, A.R.J, D.K., C.P.K and E.R. conceived the project. A.A.B., A.L.M., D.K., M.K., A.H., S.C., and N.I. performed experiments. S.P. and A.A.B. generated transgenic *Drosophila*. A.A.B., A.L.M., D.K., M.K., A.H., S.C., N.I., J.R.W. and A.R.J. analysed the data. A.L.M., A.A.B., A.R.J., J.R.W., D.K., C.P.K., E.R. and R.A.B. wrote the manuscript. C.P.K., E.R., A.R.J. and R.A.B. acquired funding.

## Conflict of interest statement

The authors declare no competing financial interests.

## Methods

### Molecular Dynamics Simulations

The *Dm*CRY-CT was excised from the protein structure, as available from the protein databank under ID 4GU5^23^, and extended by the three missing valine residues. MD simulations were then realized using the NAMD software^41,42^. The peptide, water molecules and ions were modelled using the CHARMM36 force field^43,44^; for FAD a previous parametrisation described in^45–47^ was used. Following the CHARMM force field recommendations, a 1.2 nm cutoff was used for intermolecular interactions, with the CHARMM switching function active from 1.0–1.2 nm. Pair lists were updated with distance cutoff of 1.35 nm. Periodic boundary conditions were used, and long-range Coulomb interactions were calculated using the particle mesh Ewald method. All simulations were carried out at 295 K and a pressure of 1 atm; the thermostat employed a time constant of 200 fs; the barostat was operated with a time constant of 50 fs and period of 200 fs, respectively. The histidine residue 522 was modelled as HSD (*i.e*., as neutral residue with Nδ protonated); all other ionizable residues were assigned their typical state expected for pH 7; *i.e*., lysine and arginine were present in their cationic and glutamic and aspartic acid in their anionic forms.

The peptide and/or FAD were solvated in TIP/3P water^43^ ensuring that a 25 Å layer of water surrounded the solutes in each direction. Sodium and chloride ions were added to the simulation to neutralize the simulation cell and establish a salt concentration of 50 mM (typically, this equated to 38 sodium and 30 chloride ions being present). To relax and equilibrate the systems, a protocol similar to previous studies was employed^45,46,48^. The energy was minimized and subject to MD simulations, employing progressively relaxed restraining potentials for the peptide and FAD. Specifically, the system was minimized using a steepest descent algorithm (10000 steps), while restraining the peptide and FAD atom position by a strong harmonic potential with force constant k = 18 kcal/mol/Å^2^. This was followed by an initial 5 ns MD simulation with equally restrained FAD and the peptide atom positions. A subsequent 5 ns simulation restrained only the protein backbone atom positions with *k* = 2 kcal/mol/Å^2^. The final equilibration step consisted in a 20 ns simulation with no atom position restraints. The first equilibration simulations employed an integration time step of 1 fs, while all subsequent steps used timesteps of 2 fs and constrained the hydrogen bond length to their equilibrium values using the SHAKE algorithm. Extensive production runs were run for simulation times and configurations as summarized in Table 1. A typical system, namely the *configuration I* simulations, comprised besides the peptide and FAD, 17038 H_2_O, 16 Cl^-^, 17 Na^+^, totalling to 52095 atoms, in a simulation box of dimensions 82 Å × 76 Å × 82 Å.

Snapshots were saved ever 20 ps. Data were visualized in VMD^49^ and analysed using Python scripts using the MDAnalysis^50,51^ package after reassigning the FAD position to correspond to the mirror image closest to the peptide. Interaction energies were calculated in NAMD from the saved trajectories.

### *Dm*CRY-CT and FAD in vitro binding assays

#### Peptide solubilization

##### *Dm*CRY-CT peptide

[YPERIIDLSMAVKRNMLAMKSLRNSLITPPPHCRPSNEEEVRQFFWLA] (Sigma Aldrich, UK) was solubilized in 50 mM HEPES (Fisher Scientific), pH = 7.4 or the same buffer supplemented with 1% (w/v) FOS-CHOLINE-12 n-Dodecylphosphocholine (FC-12) (Generon). The peptide solution was incubated in buffer for at least 30 min at room temperature, after which any insolubilities were removed by centrifugation at 14,000 × g for 15 min at room temperature, after which the supernatant was collected for measurement.

##### Size-exclusion chromatography

Size-exclusion chromatography (SEC) was performed in the dark with a Shimadzu UFLC Prominence HPLC System and GE Healthcare Superdex 75 increase 10/300 GL column. All solution mixtures were prepared in 50 mM HEPES (Fisher Scientific), pH = 7.4. *Dm*CRY-CT (30 μM) and FAD (30 μM) (disodium salt hydrate, Sigma-Aldrich) were run independently and benchmarked against 30 μM of the peptide, epidermicin. *Dm*CRY-CT and FAD mixtures at a range of ratios were prepared and run under red light. In addition, a mixture of *Dm*CRY-CT (30 μM) and FAD (60 μM) was run following illumination with 450 nm LED for 15 min at 0.5 mW power. The mixtures were run in the presence or absence of FC-12. The samples were filtered with 0.2 μm syringe filters (Minisart RC, Sartorius) prior to loading. Sample (100 μl) was injected on to the column and run with 0.75 ml/min flow rate. Absorbance of the elute was monitored at 280 nm, excitation performed at 450 nm, and fluorescence emission recorded at 530 nm.

##### Absorption and fluorescence spectroscopy

All solution mixtures were prepared in 50 mM HEPES (Fisher Scientific), pH = 7.4. Solution concentrations were determined from absorption spectra: FAD (450 nm, 11300 M^−1^cm^−1^) and *Dm*CRY-CT (280 nm, 6990 M^−1^cm^−1^ calculated based on the primary sequence using the Expasy’s ProtParam tool) measured using an Agilent Cary60 UV-Vis Spectrophotometer from 200 to 700 nm in a 10 mm quartz cuvette (Starna). The baseline for each spectrum was corrected to zero at 700 nm. Titration series of *Dm*CRY-CT and FAD were prepared with a constant FAD concentration (between 7.0 and 8.0 μM) with *Dm*CRY-CT concentration varying from 0 to 37.5 μM. The absorption spectra of the series for accurate FAD and *Dm*CRY-CT concentrations were measured using the Cary60 as described above. Fluorescence spectra were acquired using an Edinburgh Instruments FLS1000 fluorescence spectrophotometer. FAD was photoexcited by 450 nm light (4 nm excitation slit width) and the emission was collected from 460 nm to 700 nm (1.25 nm emission slit width) with 1 nm spectral resolution and 0.5 s/nm dwell time. The lifetime measurements were performed from the same titration series using the FLS1000 with photoexcitation at 450 nm by an Agile Supercontinuum Laser (5 nm excitation slit width) at 1 MHz frequency. Emission was followed at 530 nm (10 nm emission slit width) until 10000 counts at 50 ps time-resolution. The instrument response function (IRF) at 450 nm was measured using 50% Ludox with identical settings except for decreased beam intensity.

##### Stern-Volmer Analysis

FAD fluorescence spectra were normalised to the FAD concentration based on the absorption spectra. The fluorescence was then integrated from 460 nm to 700 nm (*I*), divided by the integrated intensity in the absence of *Dm*CRY-CT (*I*_0_), and plotted as a function of the *Dm*CRY-CT / FAD concentration ratio. The data were fitted to Eq. 1^52^.

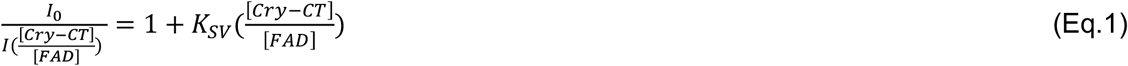

Where *K*_SV_ is the Stern-Volmer constant. For the datapoints where the molar ratio of *Dm*CRY-CT / FAD was greater than 1, the fit was performed with an unrestricted constant with a lower bound of 1.

FAF+D fluorescence lifetime data were fitted to the sum of two exponential decays (Eq. 2) as expected for FAD^38^, with iterative reconvolution using the IRF.

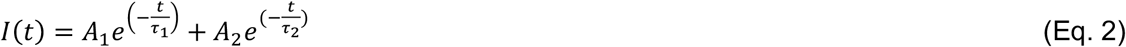

The Stern-Volmer analysis was performed with the lifetime datasets according to Eq. 3^52^.

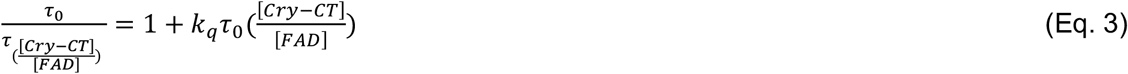

where *τ* is the fluorescence lifetime, *τ* _0_ the lifetime in the absence of *Dm*CRY-CT and *k*_q_ is the quencher rate coefficient.

### *Dm*CRY-CT and FAD fluorescence microscopy

*Dm*CRY-CT and FC-12 were the same materials used above. FAD was supplied by Thermo Scientific Chemicals. All solution mixtures were prepared in 50 mM HEPES (Dojindo), pH = 7.4. Microscopy was performed in a custom microscope (described in Ikeya and Woodward 2021^35^). Laser light of 450 nm, passed through a 100 μm diameter multimode fibre, was collimated *via* an aspheric lens (*f* = 11 mm) before focussing to generate a beam width of 17.2 μm. This beam was then transmitted to the sample via a dichroic mirror (transmission cutoff wavelength 470 nm) through a long pass excitation filter (cutoff wavelength 500 nm). Collected sample fluorescence was captured by a sCMOS (ORCA Flash 4.0 V3, Hamamatsu) imaging camera or a diode array spectrometer.

Samples of *Dm*CRY-CT (100 μM) in 50 mM HEPES buffer, pH 7.4, were prepared and stored at room temperature overnight before centrifuging (10,000rpm at 4°C for 40 minutes) to remove precipitate. A small volume (1/100th of the *Dm*CRY-CT solution volume) of FAD solution was added to adjust the solution to the required FAD concentration. When this step was required, experiments were performed in both room light and no room light (under 630nm light). Solutions were measured under the microscope within 10 minutes of preparation. For experiments performed with the addition of FC-12, experiments were performed according to two different protocols. In the first protocol, *Dm*CRY-CT solution was prepared to include 1% FC-12. In the second protocol, FAD solution was first added to the *Dm*CRY-CT solution and incubated for 8-hours to allow for particle formation. Next, 10 μL of FC-12 solution in HEPES buffer was added to 90μL of the FAD / *Dm*CRY-CT solution along with a matching control in which 10 μL of HEPES buffer was added, before both solutions were imaged under the microscopen (Extended Data Fig.4A).

Fluorescence spectra were recorded using a mini-spectrometer (C10083CA, Hamamatsu). Excitation was delivered through a 25 µm-diameter multimode optical fiber, producing a focused beam with a 4.3 µm diameter to selectively illuminate aggregates while minimizing excitation of other components. The excitation intensity at the sample was 4.15 kW/cm^2^.

### *Drosophila* stocks

Flies were maintained on standard cornmeal food at 25 °C on a 12:12 light/dark cycle until wall-climbing third instar larvae (L3) emerged. To avoid light-dependent degradation of *Dm*CRY-CT, larvae were reared in the dark throughout the day of recording. Fly stocks used were obtained from the following sources: *elaV*^*C155*^*-Gal4*; *+*; *cry*^*03*^ (Bloomington Drosophila Resource Centre #458), *w*^*-*^; *UAS-Myc-DmCRY-CT*^*R504E*^; *+, w*^*-*^; *UAS-Myc-DmCRY-CT*^*R524E*^; *+, w*^*-*^; *UAS-Myc-DmCRY-CT*^*R524A*^; *+, w*^*-*^; *UAS-Myc-DmCRY-CT*^*R524K*^; *+, w*^*-*^; *UAS-Myc-DmCRY-CT*^*K503A,R524A*^; *+, w*^*-*^; *UAS-myc-DmCRY-CT*^*V531K*^; *+, w*^*-*^; *UAS-myc-DmCRY-CT*^*V541K*^; *+* were made by the Baines group in collaboration with Manchester University’s Genomic Technologies and Fly facility.

### Cloning of myc-CT constructs

The (3xMyc)-*Dm*CRY-CT construct was made by gene synthesis (Eurofins Genomics). All subsequent variants of *Dm*CRY-CT were generated in house by the Genomic Technologies facility at the University of Manchester via site directed mutagenesis of the *Myc-DmCRY-CT* construct. Variants were assembled into the pJFRC-MUH plasmid backbone (Addgene #26213) via the 5’NotI and 3’ XbaI restriction sites. Transgenic injections were carried out by the Manchester University Fly Facility using the *y w M(eGFP,vas-int,dmRFP)ZH-2A;P*{*CaryP*}*attP40* line (stock 13-40, University of Cambridge Fly Facility) utilising the Phi31 integrase system.

### Electrophysiology

Wall-climbing third instar (L3) larvae were dissected in extracellular saline (135 mM NaCl, 5 mM KCl, 4mM MgCl_2_·6H_2_O, 2 mM CaCl_2_, 5 mM TES and 36mM Sucrose, pH 7.15) as described previously^8,26,53^, with a red filter applied to both the dissecting light and compound microscope to minimize *Dm*CRY degradation during dissection and recordings. Thick-walled borosilicate glass electrodes (GC100F-10; Harvard Apparatus) were fire-polished to a resistance of 10-15 MΩ and back-filled with intracellular saline (140 mM K^+^-D-gluconate, 2 mM MgCl_2_·H_2_O, 2 mM EGTA, 5 mM KCl and 20 mM HEPES, pH 7.4). KCl and CaCl_2_ were obtained from Thermo Fisher Scientific, sucrose from BDH, and the remaining chemicals from Sigma-Aldrich. Recordings were performed using either an Axon Multiclamp 700B amplifier & Digidata 1550A analogue-to-digital converter, or an Axopatch 200B & Digidata 1440A (Molecular Devices). We targeted the easily identifiable aCC motoneuron for recordings^8,26,53^. Only cells with an input resistance of >500 MΩ were accepted. Mecamylamine (1 mM) was applied to preparations to synaptically isolate aCC from endogenous excitatory cholinergic input. Cells were recorded in current clamp mode with a small amount of depolarizing current injected to provide stable action potential firing (∼ 5 -7 Hz, the aCC motoneuron is quiescent when isolated from its synaptic drive).

### Photoactivation and magnetic field exposure

BL exposure was applied using an LED (470 nm; ThorLabs or Cairn) at a power of 2.2 mW.cm^-2^, which has previously been used for *Dm*CRY photoactivation^8,54,55^. Following establishment of a stable baseline of action potential firing, aCC cells were recorded for 20 s, followed by 30 s of BL exposure. Magnetic field exposure was supplied by two NdFeB static magnets mounted around the preparation to provide a 100 mT (±5 mT) magnetic field. Field strength was measured using a 5180 Gauss/Tesla Meter (F.W. Bell): see Bradlaugh *et al*., 2023^8^ for more details.

### Statistical Analysis of Electrophysiology Data

The experimenter was blinded to genotype throughout recordings and data analysis. A sample size of 8-10 cells per genotype, per condition was used based on previously published work^8,26^, with each cell recorded being from an independent larva (i.e. only one cell per animal was recorded).

Data was tested for normality using a D’Agostino-Pearson analysis. BL sensitivity was determined using a paired t-test to compare the number of action potentials fired in the 15 s following BL on compared to 15 s prior. This was used to calculate a firing fold change (FFon/FFoff). For comparison between BL and BL+MF, the firing fold change under each condition was compared. Statistical significance for BL+MF against BL alone was tested using an unpaired two-tailed t-test. Data are shown as mean ± SEM.

## Notes

https://doi.org/10.7910/DVN/GLW1UV

### Competing Interest Statement

The authors have declared no competing interest.

