## Extended Data for "How the *Drosophila* Cryptochrome C-terminus mediates magnetosensitivity"

Extended Data Figure 1

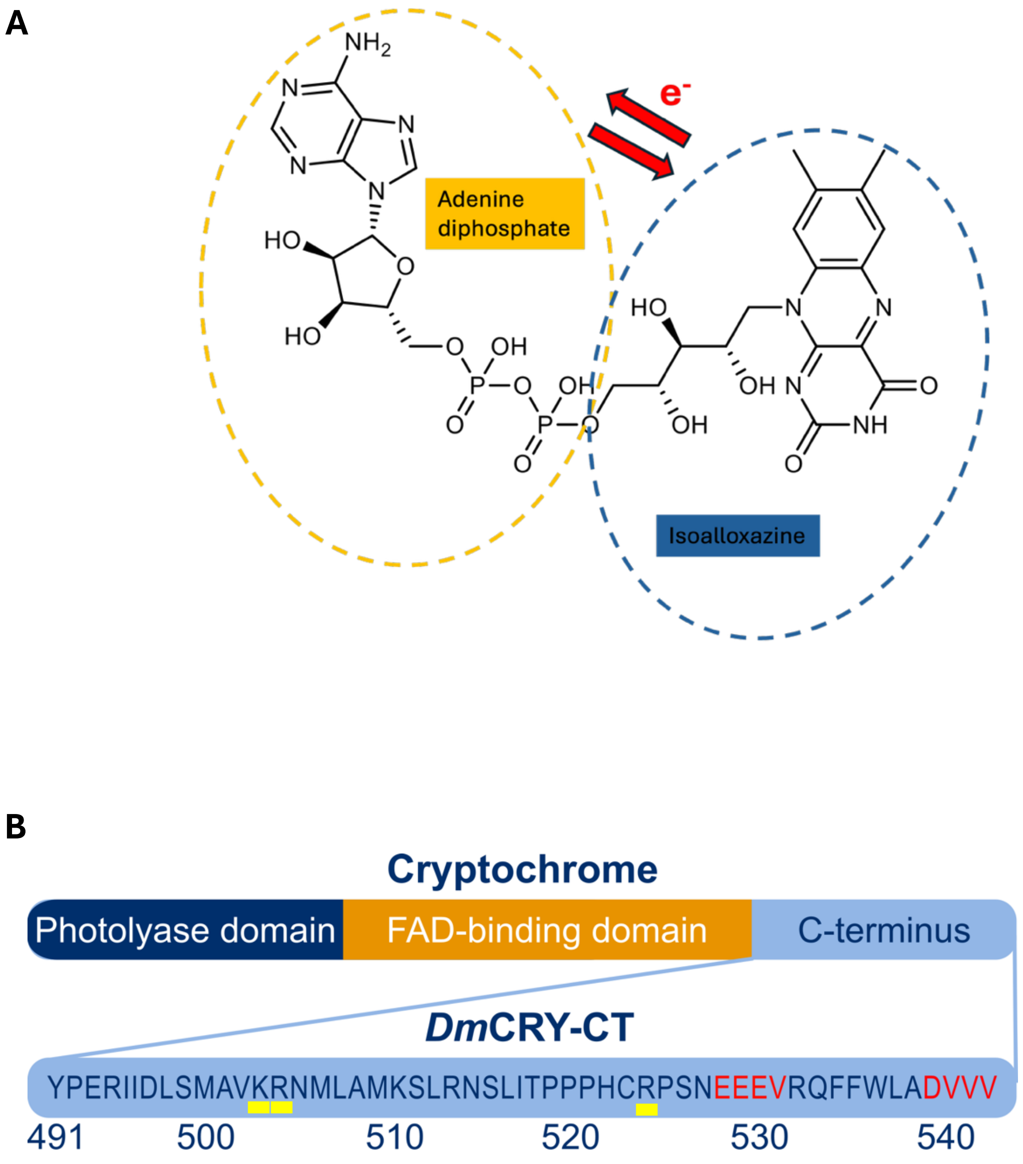


**Extended Data Figure 1. Structures of FAD and *Dm*CRY.**

A) The molecular structure of flavin adenine dinucleotide (FAD). Dashed yellow oval shows the adenine diphosphate side group, the blue dashed oval highlights the chromophore region. The intramolecular electron transfer sensitive to a MF occurs between these two groups. B) The overall structure of *Drosophila* Cryptochrome and its C-terminus, red residues denote PDZ domains, residues underlined in yellow are those predicted by MDS to be integral to binding FAD.

**MDS of FAD binding mutants.**

We investigated the effects of mutating key residues at the FAD binding sites to explore their impact on FAD binding and derive testable hypothesis. MDS were performed for the mutant structures, initiating the simulations from the bound structures and focussing on the same parameters as the wild-type configurations. In all cases, the overall fold remained intact, but the FAD binding affinity was significantly altered depending on the mutations.

For *configuration I*, we examined R504E. As expected, the introduction of a negatively charged Glu residue weakened FAD binding (Main Text Table 2). The FAD shifted from its original site, resulting in weaker interactions with markedly increased interaction energies of -110, -120, and -70 kcal/mol, respectively. The R504E mutation led to clear FAD detachment, a closure of FAD structure, and eventual binding in the groove between α-helices (Below, Extended Data Fig. 2). In the other mutants, FAD migrated to nearby residues, *e.g.*, Y491 or R513, in part folding into a closed configuration.

For *configuration II*, R524A and R524E showed increased interaction energies, reflecting weaker binding due to the replacement of the positively charged arginine with either neutral (alanine) or negatively charged (glutamic acid) residues. However, the mutations did not necessarily induce dissociation or marked site migration; in two instances the adjacent R513 maintained bound FAD, while in one a shift to R504 of *configuration I* occurred. We note that the preservation of (partial) binding observed here applies to FAD initially in a bound configuration. With E524 present, however, this configuration might not have formed from free FAD, as R513-E524 will have formed a salt bridge, rendering the interaction of FAD with R513 less forcing. The positive control mutation, R524K (positive replacing positive residue), again successfully preserved the original FAD binding. These findings highlight the importance of electrostatic interactions in FAD binding to the *Dm*CRY-CT and suggest that specific mutations may significantly alter binding behaviour, providing valuable insights for further experimental validation.

Extended Data Figure 2

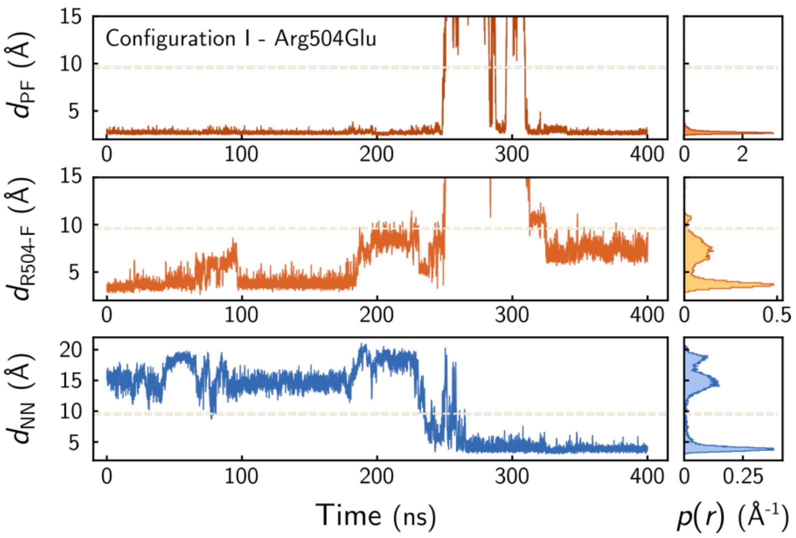


**Extended Data Figure 2. Representative time evolution of R504E-FAD binding.**

Time evolution of the peptide-FAD distance $d_{PF}$ (red; top), the distance of FAD from R504 (orange; middle), and the inter-moity distance $d_{NN}$ in FAD (blue; bottom) for FAD initially bound to the R504E mutant in *configuration I*. The FAD detaches from the binding site, is folded closed, and is recaptured by the peptide, ending up weakly bound between the two α-helices.

### Extended Data Figure 3

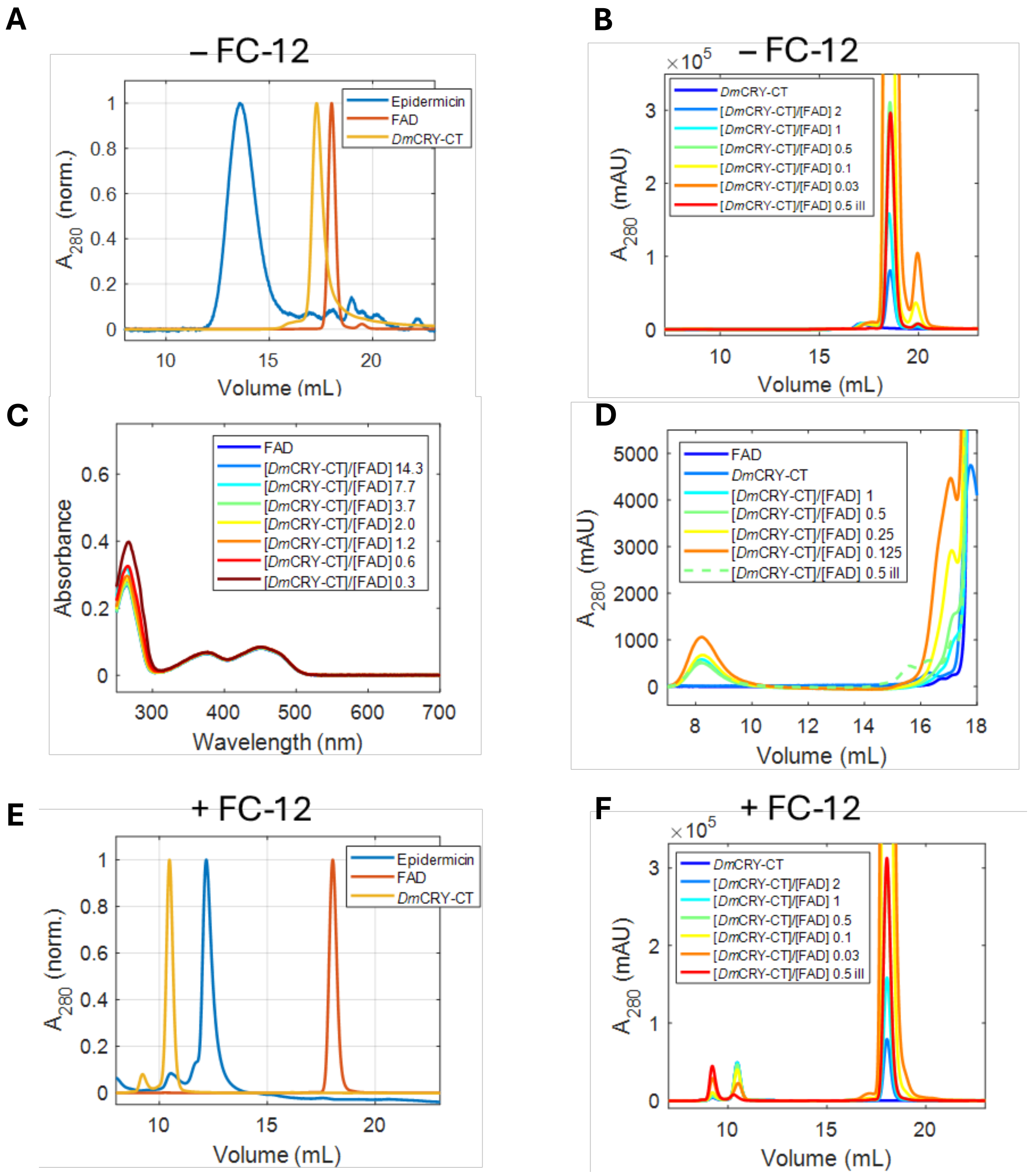


**Extended Data Figure 3. Size Exclusion Chromatography and UV-vis.**

A) The SEC elution profile of epidermicin (6.1 kDa), *Dm*CRY-CT (5.8 kDa), and FAD (0.8 kDa) run separately in the absence of FC-12. The epidermicin peptide was used as a size control to confirm that *Dm*CRY-CT elutes as a monomer, like the slightly larger epidermicin. B) The full elution profile of the samples in Fig. 2A. including the illuminated sample (ill) from Fig. 2C. C) UV-Vis spectra of *Dm*CRY-CT + FAD titration series (Fig. 2B) supernatant after centrifugation. The scattering particles are removed by centrifugation, and the FAD concentration remains almost unchanged. D) SEC elution profile detected by absorbance at 280 nm of samples prepared in ambient room light has pronounced features in comparison to dark-run samples in Fig. 2A. E) The SEC elution profile of epidermicin, *Dm*CRY-CT, and FAD run separately in the presence of FC-12. The elution of the *Dm*CRY-CT changes significantly relative to FAD and epidermicin, presumably owing to differences in hydrophobicity and thus association of FC-12 to the *Dm*CRY-CT peptide forming larger species. F) The full elution profile of the samples in Fig. 2D and illuminated (ill.) from Fig. 2F.

**Synthesis of epidermicin.**

Epidermicin was assembled using a Liberty Blue microwave peptide synthesizer (CEM), Fmoc/tBu solid-phase synthesis protocols and DIC/OxymaPure as coupling reagents. Formyl Methionine (Merck) and Pseudoproline dipeptides (Fmoc-L-Ala-L-Thr[psi(Me,Me)Pro]-OH, Fmoc-L-Val-L-Ser[psi(Me,Me)Pro]-OH, Fmoc-L-Gly-L-Thr[psi(Me,Me)Pro]-OH, Iris Biotech) were utilized to aid peptide synthesis. Cleavage and deprotection (90% TFA, 5% TIS, 5% water) yielded the crude peptide, which was precipitated using diethyl ether and lyophilized.

The resulting lyophilized peptide was purified by preparative reversed-phase high-performance liquid chromatography (RP-HPLC) and analysed by analytical RP-HPLC and liquid chromatography mass-spectrometry (LC-MS). RP-HPLC purification was performed on a Waters, XSelect, CSH, C18 column (PN: 186005493, 30 mm × 250 mm, 130 A, 5 $\mu$m). Preparative runs were carried out at 40 mL/min using a 0−40% buffer B gradient over 40 min (buffer A is 95% and buffer B is 5% aqueous $CH_3CN$, each containing 0.1% TFA). Fractions containing purified peptide were pooled and lyophilized to yield Epidermicin, *Formyl-MAAFMKLIQFLATKGQKYVSLAWKHKGTIL KWINAGQSFEWIYKQIKKLWA-amide*, mass = 31 mg, Mw = 6,071.3550 g $mol^{-1}$, n = 5.1 x $10^{-3}$ mmol, % Yield = 5.1 %. Analytical runs were performed at 1 mL $min^{-1}$ using a 0−100% buffer B gradient (over either 20 or 50 min) on a Waters, XSelect, CSH, C18 column (PN: 186005291, 4.6 mm × 250 mm, 130 A, 5 $\mu$m), with detection at 214 nm. LCMS analyses were performed on a Thermo Scientific Q-Exactive system (equipped with a HESI probe), using a Phenomenex Synergi C12 column (PN: 00F-4337-E0, 4.6 mm × 150 mm, 80 A, 4 $\mu$m). With 20 min gradient, epidermicin eluted at 13.2 min with 91.8 % purity, and 50 min gradient at 25.4 min and 90.5 % purity.

LCMS runs utilized a 0−100% buffer B gradient over 20 min at 0.8 mL/min with detection at 214 nm (buffer A, 95% and buffer B, 5% aqueous $CH_3CN$, each containing 0.1% formic acid). High resolution LCMS, m/z ($ESI^+$), reported as m/z observed / Da (assignment = m/z expected / Da): 1,214.4783 Da (1/5$[M+5H]^+$ = 1,214.4760 Da), 1,012.2277 Da (1/6$[M+6H]^+$ = 1,012.2313 Da), 867.7699 Da (1/7$[M+7H]^+$ = 867.7708 Da), 759.4245 Da (1/8$[M+8H]^+$ = 759.4254 Da), 675.1563 Da (1/9$[M+9H]^+$ = 675.1568 Da), 607.7406 Da (1/10$[M+10H]^+$ = 607.7419 Da).

Extended Data Figure 4

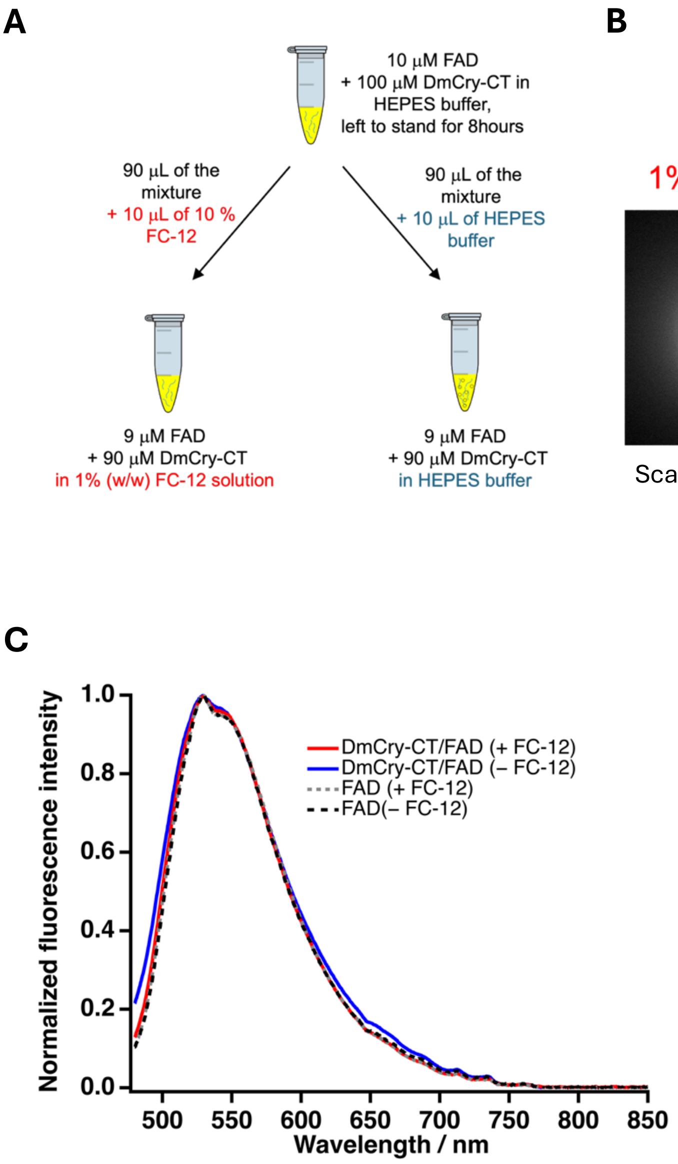


**Extended Data Figure 4. Measurements to confirm dissolution of clusters in FC12 detergent.**

A) Schematic of sample preparation. A sample is initially prepared without FC12 to allow cluster formation. The sample is then divided in two aliquots, only one of which has FC12 added, B) Fluorescence imaging of these two aliquots which reveals that no clusters are observable in the aliquot containing FC12, while the detergent free sample shows clear cluster formation. C) Fluorescence spectra of these two aliquots. In the aliquot to which FC12 was added, the shape of the fluorescence spectrum approaches that of the isolated FAD solution.

Extended Data Figure 5

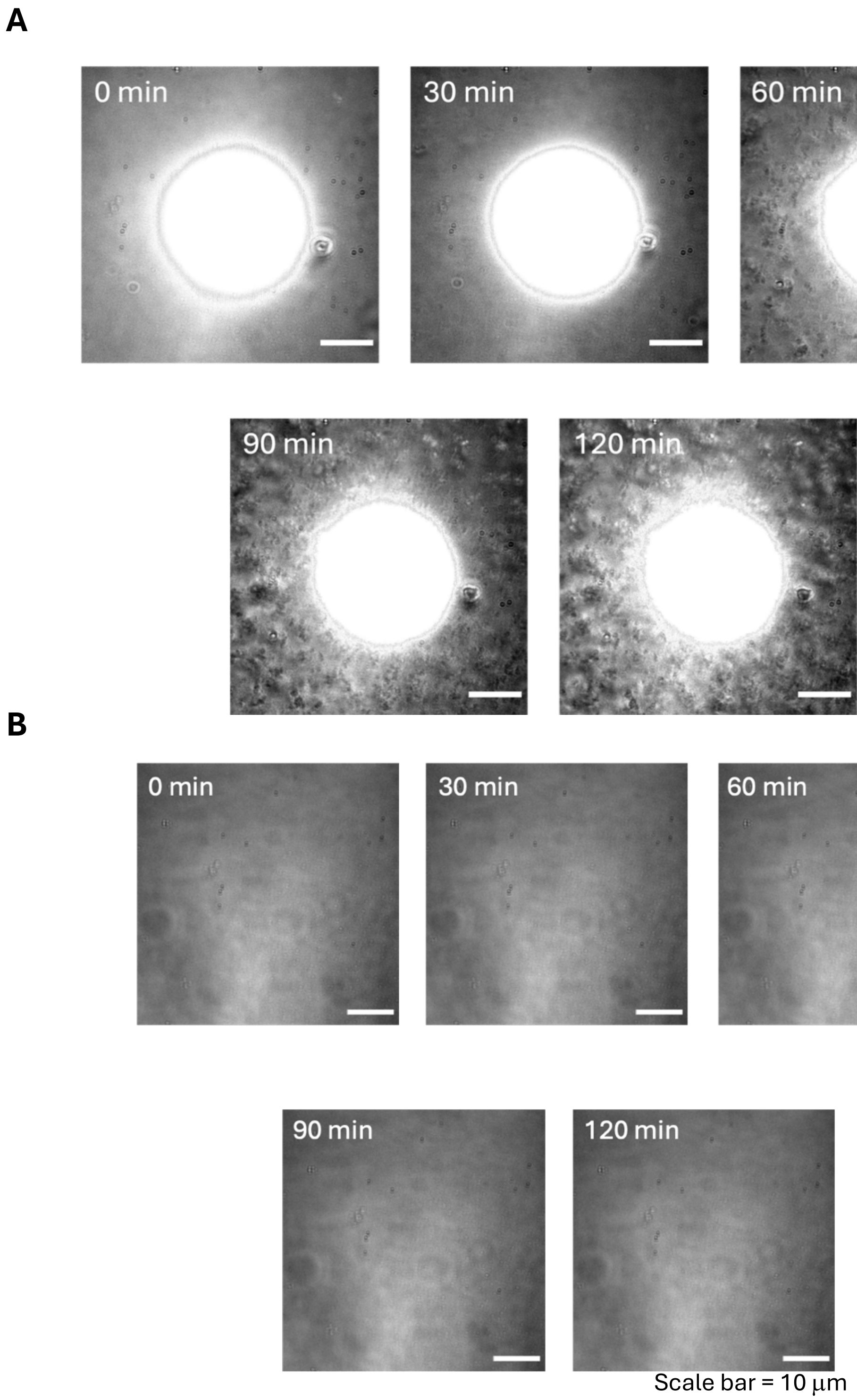


**Extended Data Figure 5. Time-lapse bright-field microscopy showing particle formation around irradiation spot.**

A) Time-lapse bright-field images for 100 μM Cry-CT and 10 μM FAD solution (pH 7.4) in the absence of FC12 under 450 nm photoexcitation. The bright field imaging mode reveals the gradual accumulation of particles in the area surrounding the irradiation spot during photoexcitation of the sample. This data is taken from the same data set presented in Fig. 3C with the contrast adjusted to expose the bright field signal in the region surrounding the focussed laser irradiation spot. B) Time lapse of bright field microscope images for a non-irradiated control experiment (450 nm laser off). Video available at Harvard Dataverse: https://doi.org/10.7910/DVN/GLW1UV.

Extended Data Figure 6

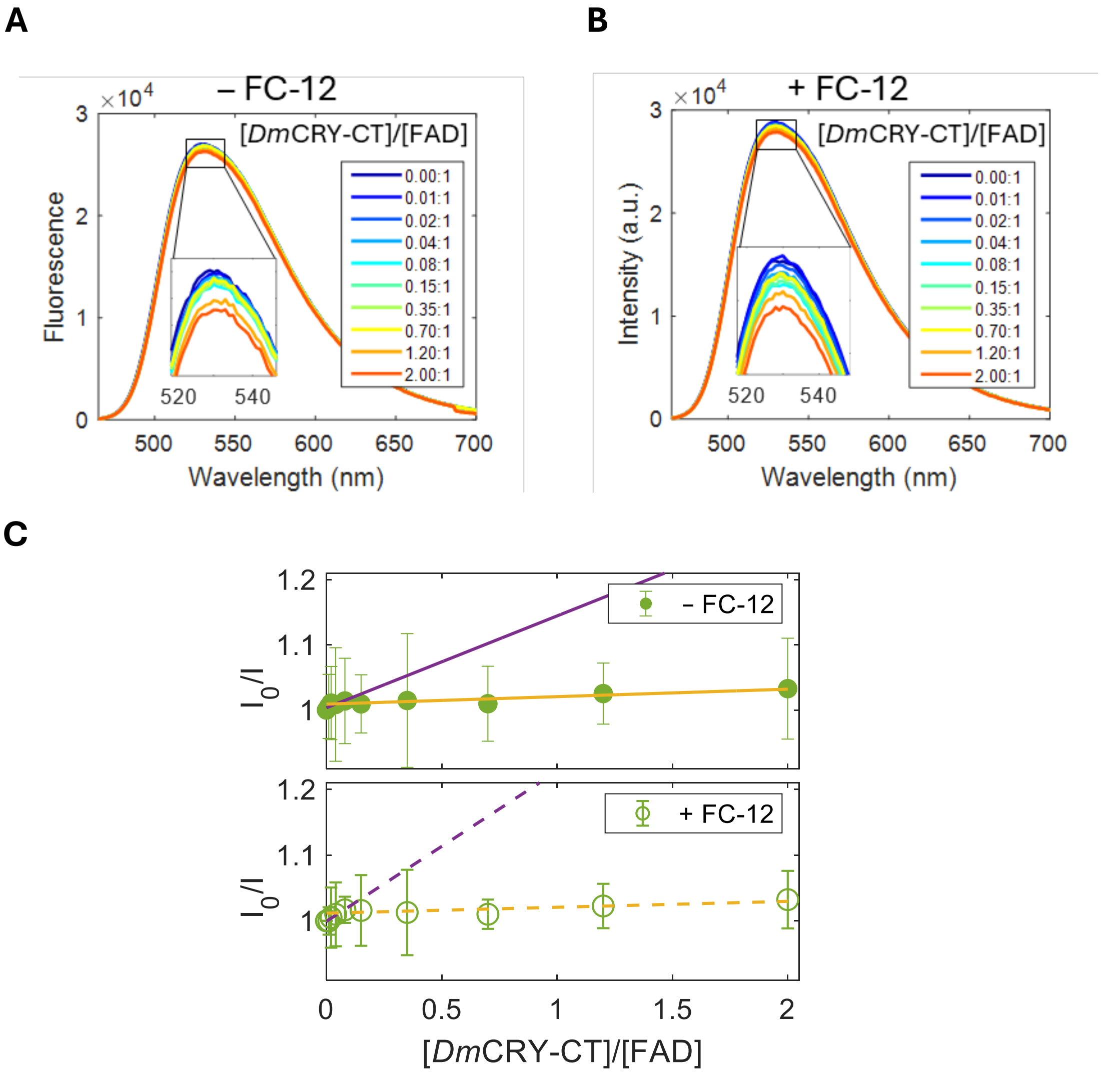


**Extended Data Figure 6. Fluorescence spectra.**

A-B) Fluorescence spectra of FAD at constant concentration (7.4 μM) with increasing [*Dm*CRY-CT] / [FAD] ratios (see in-panel legends) and in the absence (A) and presence (B) of FC-12 demonstrate a small but significant and reproducible quenching of the FAD fluorescence upon addition of *Dm*CRY-CT. The lack of difference to the quenching ± FC-12 suggests negligible quenching of fluorescence from FAD that 'seeds' the formation of oligomers / aggregates. This is consistent with FAD fluorescence measured from peptide particles (Fig. 3D). C) The Stern-Volmer analyses of the fluorescence data as a function of [*Dm*CRY-CT] / [FAD] in the absence (top) and presence (bottom) of FC-12: $I_0$ is the fully integrated fluorescence intensity from the spectrum of FAD alone; $I$ is the integrated fluorescence intensity from each spectrum at different [*Dm*CRY-CT] / [FAD] ratios; and the error bars are the standard deviation of three replicates. The purple lines (solid without FC12, dashed with) is the single polynomial fit from 0 to 0.1 *Dm*CRY-CT/FAD concentration ratio, and the yellow lines from 0.1 to 2. This reveals that quenching increases to a ratio of ~0.1, and then saturates remaining unchanged thereafter, even when the peptide is in excess. The trend is the same both ± FC-12. These data are consistent with quenching being predominantly from a small, soluble sub-population of complexes that don't form oligomers or aggregates.

### Extended Data Figure 7

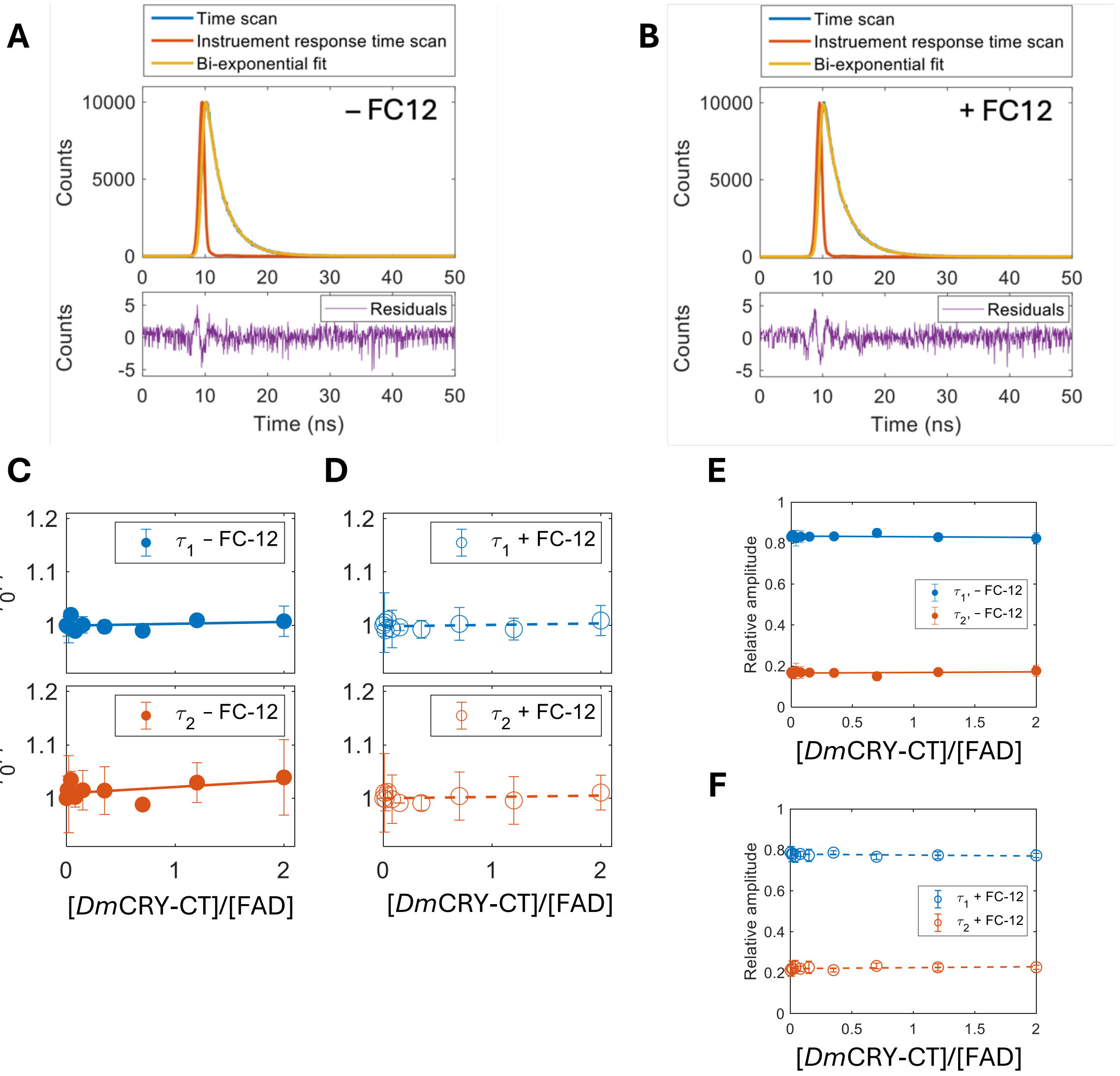


**Extended Data Figure 7. Fluorescence lifetimes.**

Fluorescence lifetimes of FAD following excitation at 450 nm at increasing [*Dm*CRY-CT] / [FAD] ratios, ± FC-12. A-B) The time scan, instrument response function (450 nm), bi-exponential fit and the residuals (see in-panel legend) for the lifetimes of FAD alone, in the absence (A) and presence (B) of FC-12. C-D) Stern-Volmer analyses for the two FAD fluorescence lifetimes, 2.4 ns ($\tau_1$) and 4.1 ns ($\tau_2$), as a function of [*Dm*CRY-CT] / [FAD] in the absence (C) and presence (D) of FC-12. $\tau_0$ is the lifetime of FAD alone; $\tau$ is the lifetime at each different [*Dm*CRY-CT] / [FAD] ratio; and the error bars are the standard deviation of three replicates. If the observed quenching of FAD fluorescence spectra in Extended Data Figure 6 originated from an increase in the kinetics of competing processes, one would expect a change in fluorescence lifetimes as a function of increasing *Dm*CRY-CT concentration. The data in this figure show this is not the case. The two fluorescence lifetimes for FAD are consistent with those previously published[37], with one corresponding to its 'open' confirmation and the other to the 'closed'. For FAD solutions both ± FC-12, neither lifetime changes upon addition of the peptide. This is consistent with the observed quenching originating from formation of a small population of non-emissive complexes that don't form aggregates. These complexes are therefore likely to be a result of direct association between FAD and the *Dm*CRY-CT. E-F) The relative amplitude of the two FAD lifetimes in the absence (E) and presence F) FC-12 doesn't vary across the titration series, suggesting that complex formation has a negligible impact on the relative proportions of the 'open' and 'closed' forms of FAD.

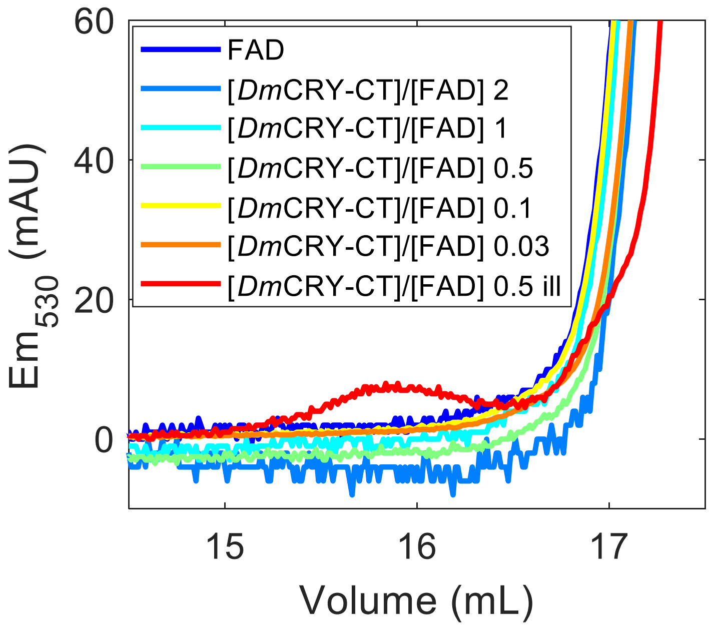


**Extended Data Figure 8. SEC Emission.**

An elution peak is observed at 15.9 mL in the illuminated sample of [*Dm*CRY-CT] / [FAD] ratio = 0.5 (red) when the elute is excited at 450 nm and emission detected at 530 nm. This corresponds to the additional peak observed at the same elution volume by absorption at 280 nm upon illumination (Fig. 2C). Emission at 530 nm is from FAD, and confirms that it is bound to the small, soluble complexes, but this only becomes visible when in sufficient quantities and requires illumination.

Extended Data Figure 9

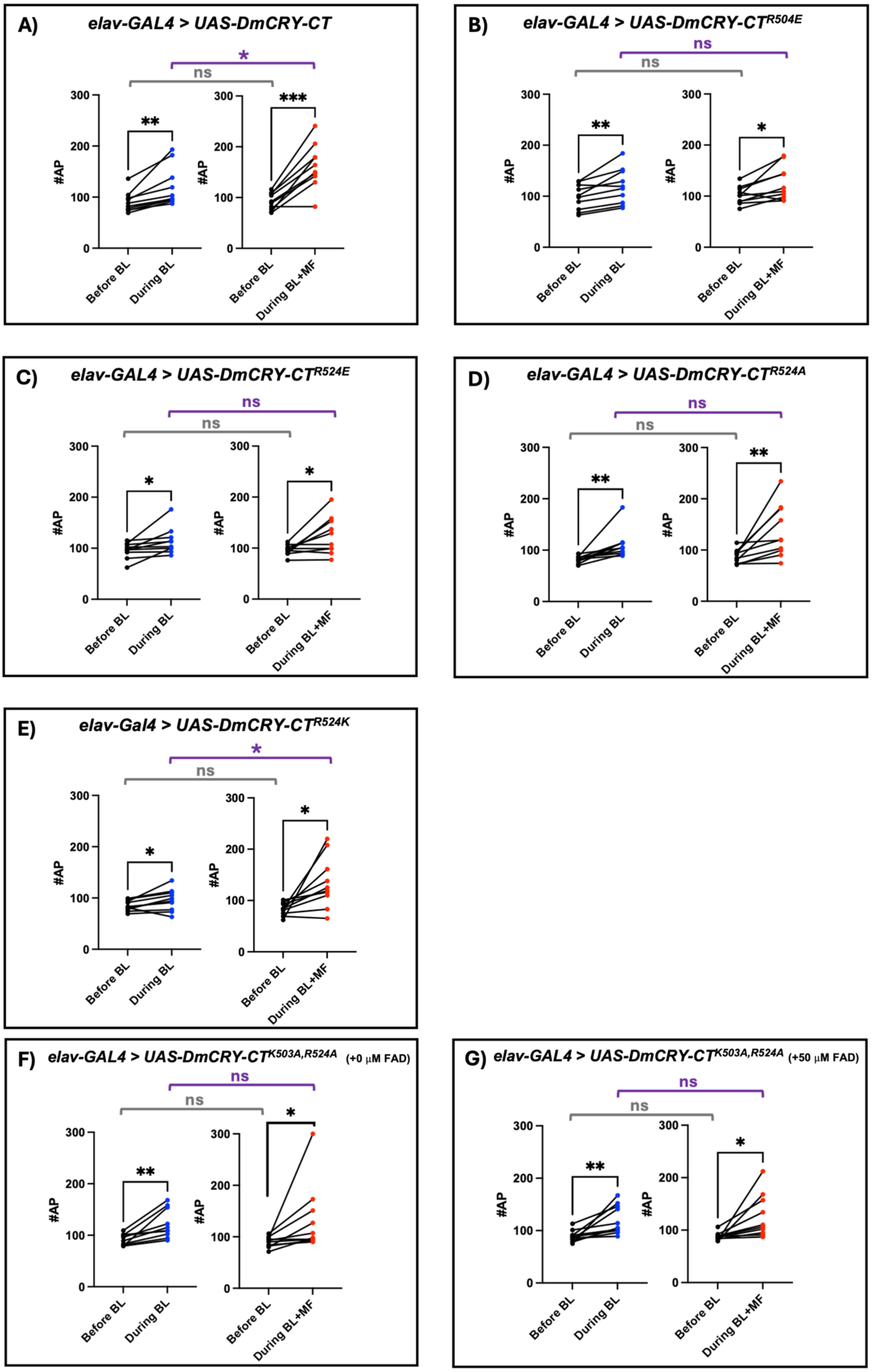

**Extended Data Figure 9. Supporting electrophysiological data for FAD binding mutants.**

A-E). Raw AP counts for each neuronal recording in both BL (blue data points) and BL+MF (red data points) for *UAS-DmCRY-CT* transgenics with point mutations designed to alter FAD binding. Number of APs in the 15 s preceding exposure *vs* number of APs in 15 s during exposure. Paired t-tests(two-tailed) show consistent BL responses across all variants. Unpaired t-tests (two-tailed) on raw AP counts between 'before' exposures were compared between BL and BL+MF (grey line) and for during exposure (purple line). Similarly to the Firing-Fold data presented in the main text, a significant effect of the MF is only seen for (A) wild-type *Dm*CRY-CT, and for the (E) UAS-*Dm*CRY-CT$^{R524K}$ transgenic in which a positive residue remains at aa:524, facilitating FAD binding. F) A double substitution of a positive for neutral residue at both binding configurations (K503A, R524A) still supports BL sensitivity but does not support MF potentiation. G) Supplementary FAD (50 μM) provided through the patch pipette did not rescue the reduced FAD binding affinity and no MF potentiation was observed. Data are derived from independent animals, one recording per animal. n=10 for each genotype in both BL and BL+MF. Error bars denote ±SEM, ns *P* ≥ 0.06, * *P* ≤ 0.05, ** *P* ≤ 0.01, *** *P* ≤ 0.001, **** *P* ≤ 0.0001.

Extended Data Figure 10

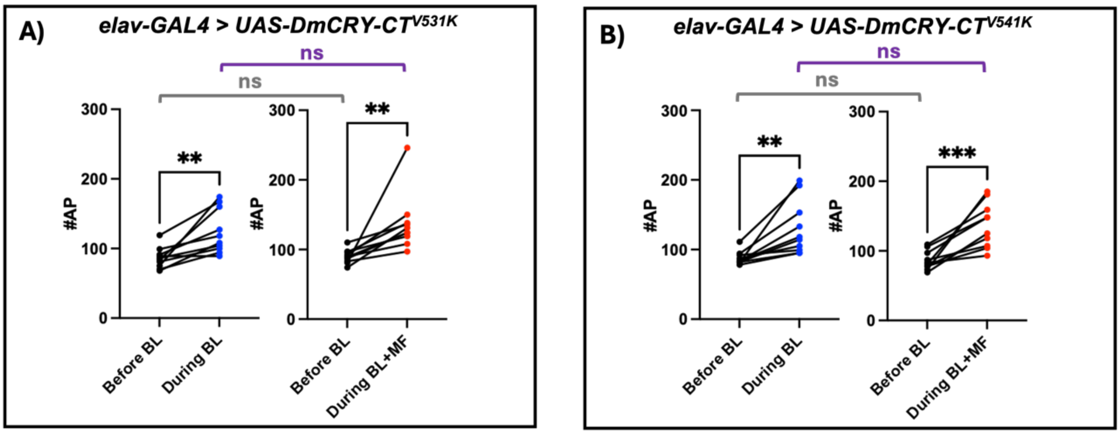


**Extended Data Figure 10. Supporting electrophysiological data for PDZ domain mutants.**

A-B). Raw AP counts for each neuronal recording in both BL (blue data points) and BL+MF (red data points) for *UAS-DmCRY-CT* transgenics with point mutations affecting two PDZ domains. Number of APs in the 15 s preceding exposure *vs* number of APs in 15 s during exposure. Paired t-tests(two-tailed) show consistent BL responses in both variants. Unpaired t-tests (two-tailed) on raw AP counts between the 'before' exposures revealed no significant difference (grey line), and no significant potentiation when comparing responses during exposure, BL *vs* BL+MF (purple line). Data are derived from independent animals, one recording per animal. n=10 for each genotype in both BL and BL+MF. ns $P \geq 0.06$, * $P \leq 0.05$, ** $P \leq 0.01$, *** $P \leq 0.001$, **** $P \leq 0.0001$.

Extended Data Figure 11

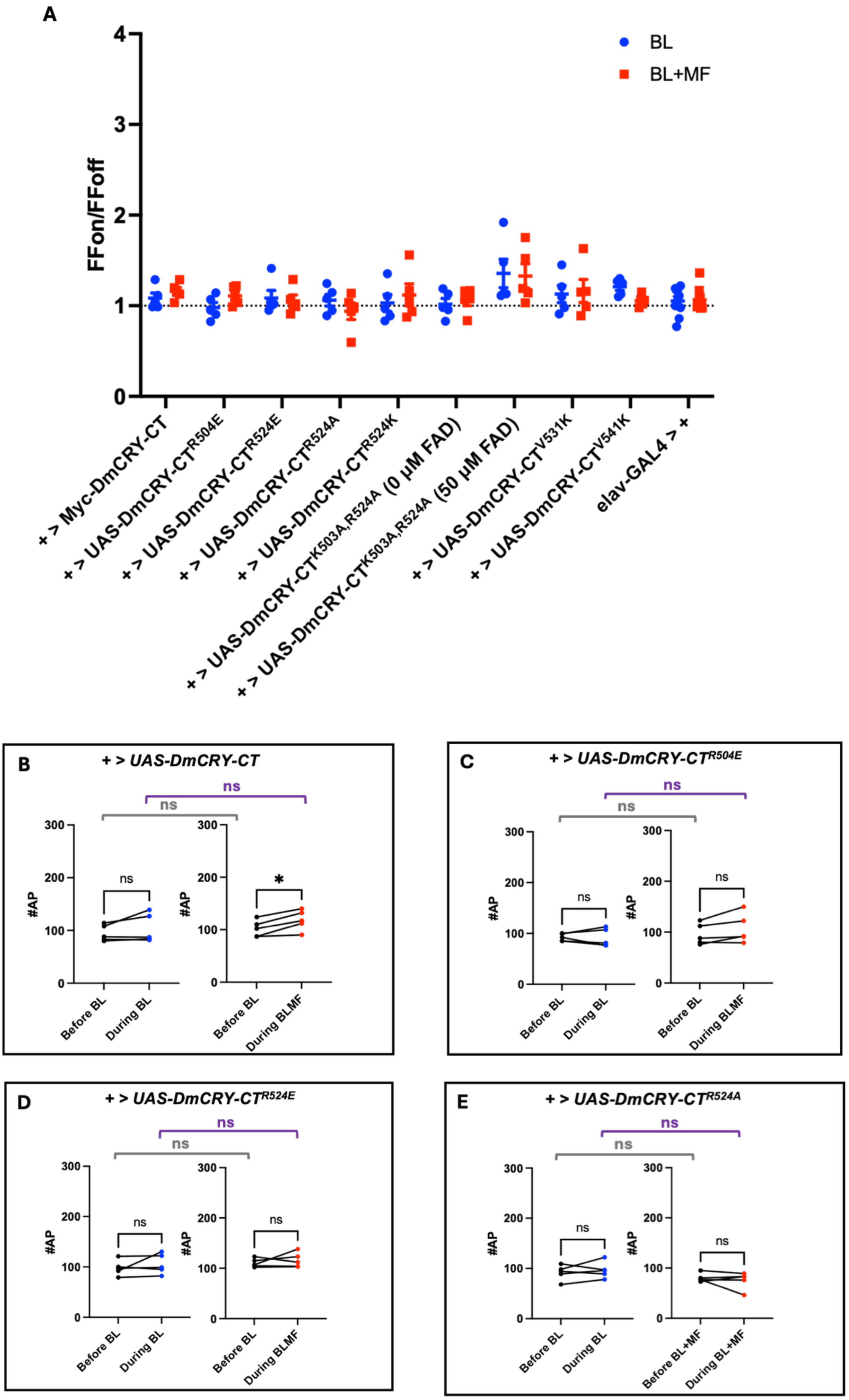

### Extended Data Figure 11

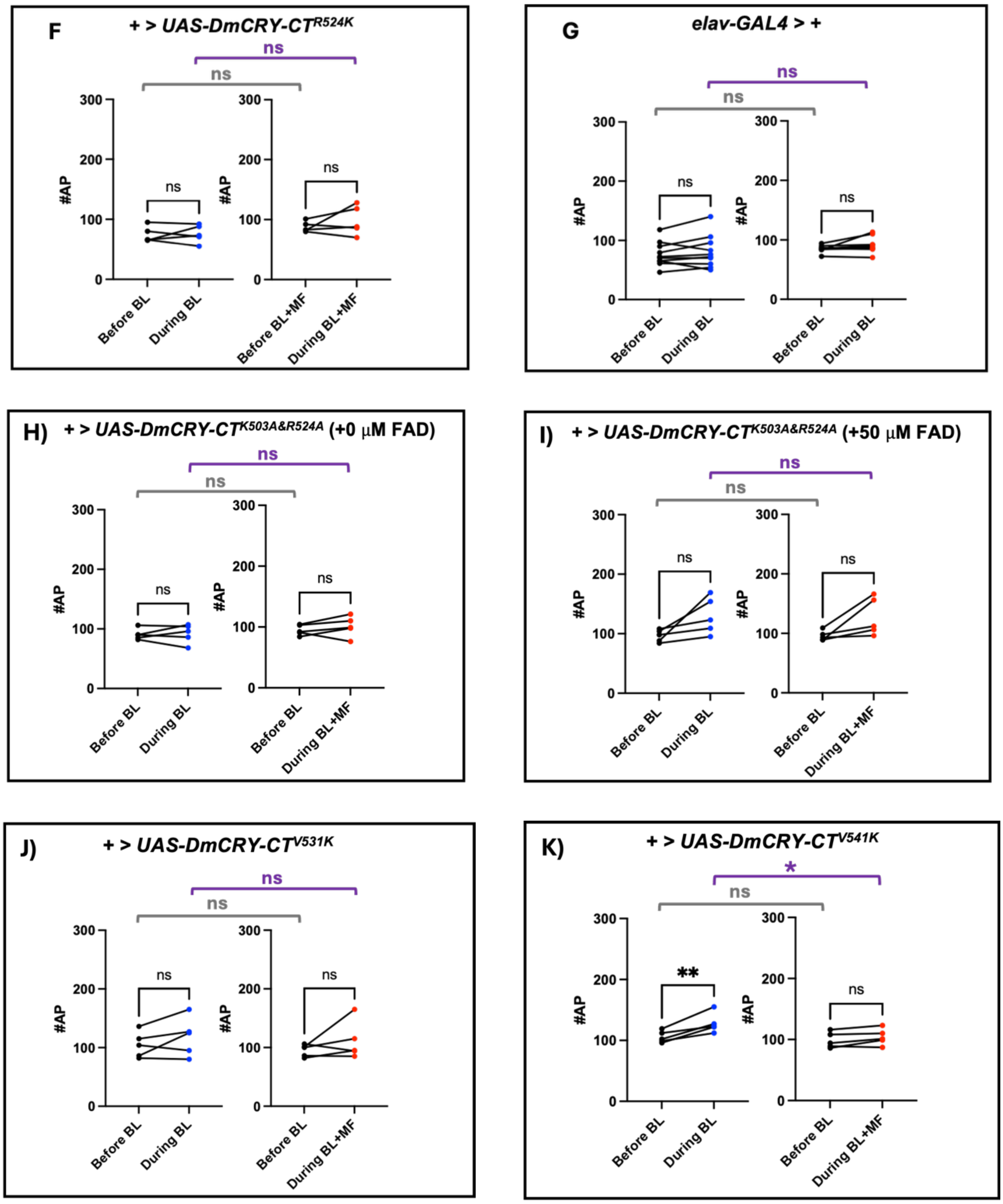

**Extended Data Figure 11. Supporting electrophysiological data for genetic controls.**

A). Averaged data for the binary expression elements of *GAL4* and *UAS* when not combined, without the GAL4 and UAS components combined in a single fly the transgenic element (UAS-) is not expressed and no significant BL or BL+MF response is seen. 2-Way ANOVA of controls in both BL and BL+MF revealed no significant interaction ($F_{(9,90)} = 0.609$, $P = 0.7987$). For the *elav-GAL4; ; Dmcry$^{03}$* driver line n=10 for both BL or BL+MF, n for each respective *UAS-DmCRY-CT* transgenic genotype =5 for both BL or BL+MF. Dashed grey line represents no change in firing fold frequency. B-G). Raw AP counts for each neuronal recording in both BL (blue data points) and BL+MF (red data points) for constituent (and separate) *elav-GAL4* and *UAS-DmCRY-CT* transgenics. Number of APs in the 15 s preceding exposure *vs* number of APs in 15 s during exposure. Paired t-tests(two-tailed) show no consistent BL or BL+MF response without the expression of a *Dm*CRY-CT transgenic. H-I). FAD supplementation did not have a significant effect on BL or BL+MF sensitivity without expression of the transgene but a trend towards greater light sensitivity in cells supplemented with FAD (50 μM) may be present. J-K). Raw AP counts from neuronal recordings of non-expressed PDZ mutants show a similar BL and BL+MF insensitivity with the possible exception of (K) + > *UAS-DmCRY-CT$^{V541K}$* which shows slight light sensitivity, possibly originating from a ‘leaky’ UAS. Unpaired t-tests (two-tailed) on raw AP counts between ‘before’ exposures were compared between BL and BL+MF (grey line) and for during respective exposures (purple line). Data are derived from independent animals, one recording per animal. Error bars denote ±SEM, ns $P \geq 0.06$, * $P \leq 0.05$, ** $P \leq 0.01$, *** $P \leq 0.001$, **** $P \leq 0.0001$.
